# Structure of active methyl-CoM reductase, Earth’s main methane producer

**DOI:** 10.1101/2025.04.26.650772

**Authors:** Christopher J. Ohmer, Macon J. Abernathy, Darya Marchany-Rivera, David G. Villareal, Philipp S. Simon, Artem Lyubimov, Asmit Bhowmick, Kuntal Chatterjee, Hiroki Makita, Vandana Tiwari, Medhanjali Dasgupta, Stephen Keable, David Mittan-Moreau, Daniel W. Paley, Nigel W. Moriarty, Humberto Sanchez, Daniel J. Rosenberg, Leland B. Gee, Roberto Alonso-Mori, Nicholas K. Sauter, Aaron S. Brewster, Uwe Bergmann, Vittal K. Yachandra, Aina E. Cohen, Junko Yano, Jan F. Kern, Ritimukta Sarangi, Stephen W. Ragsdale

## Abstract

Our work reveals the structure of the active state of Methyl-Coenzyme M Reductase (MCR), the key and rate-limiting enzyme in biological methane formation. We find large differences between the active Ni(I) and inactive Ni(II) proteins and provide insight into how nature makes and breaks the C-H bond of methane. The Ni(II)-F430 center in inactive MCR contains four planar nitrogen ligands, a lower axial glutamine oxo, and an upper axial thiolate. The Ni(I)-enzyme replaces the axial ligands with a single water. The one-electron redox change results in movement of the Ni ion and upward swing of the β-lactam ring in the tetrapyrrole coupled to a domino-like protein quake through second sphere residues, inter-subunit interactions, a substrate tunnel, affecting even the dimensions of the unit cell. These structural changes lead Ni(I)-MCR to release a charge clamp that, in the Ni(II) state, locks down substrate Coenzyme B. Determining the Ni(I)-MCR structure required development of rigorous anaerobic crystallographic techniques. Validation of the MCR redox state was accomplished by in-line and parallel spectroscopic and unit cell analyses. This structure has large implications for developing technologies to limit methane emissions and efficiently produce biofuels. Methodology described here will enhance structural biology for other oxygen-sensitive enzymes.

## Introduction

Methane played a vital role in early Earth, forming a heat-trapping greenhouse haze that warmed our planet enough to allow for the emergence of life^1,2^. Almost all the methane in our atmosphere is produced by methanogenic archaea, which emit 1 gigaton of methane annually, through the activity of Methyl-Coenzyme M Reductase (MCR), an extremely oxygen-sensitive nickel enzyme^3^. Furthermore, as the central component of natural gas, it is the second most utilized energy source in the U.S. due to its high energy output (56 kJg^-1^), low cost, and small carbon footprint compared to coal^4^. However, methane is a Janus-faced molecule, since, as an increasingly concerning greenhouse gas, it is 81-fold more potent than CO_2_ over 20 years^5^, and the estimated social cost of methane emissions (biotic and anthropogenic sources) is 50-100 times greater than that of CO_2_ ^6^. In the lower atmosphere, most emitted methane is removed by aerobic and anaerobic methanotrophic microbes^7^. Once methane escapes into the upper atmosphere, it has a short lifetime of about a decade due to its removal by reaction with hydroxyl and chlorine radicals.^8^ Understanding Nature’s process of activating the inert C-H bond and releasing methane to the lower atmosphere will play a vital role in developing methods to mitigate methane emissions.

The final step in methane synthesis, and the first step in methane oxidation, is carried out by Methyl Coenzyme M Reductase (MCR). MCR uses a Ni-tetrapyrrole, the hydrocorphin F_430,_ at its active site to catalyze the conversion of methyl-coenzyme M (CH_3_SCoM) and Coenzyme B (HSCoB) to methane and a mixed disulfide (CoBSSCoM) (Ext. Fig 1). The first crystal structures of MCR, solved by Ermler *et.al.*, were of the Ni(II) state and revealed the 300 kDa enzyme as a hexamer in a ⍺_2_β_2_ɣ_2_ configuration with two active sites^9^. In the EPR silent MCR_ox1-silent_ structure, the Ni(II) coordinates to the four planar nitrogen atoms of F_430_, with two additional axial ligands: the thiolate of inhibitor HSCoM and an oxo group from a Gln residue (Ext. Fig. 1). A wide substrate channel extends ∼50 Å into the active site pocket, with HSCoB anchored near the protein surface by ionic interactions between its anionic phosphate group and three positively charged residues (Arg, methyl-His, and Lys). The mercaptoheptanoyl-threonine group of HSCoB threads down the channel with the thiol group pointed toward the Ni of F_430_, lying at the bottom of the tunnel. The other structure presented by Ermler *et.al.*^9^ was of the Ni(II)-MCR_silent_ state containing bound product CoBSSCoM and was similar to MCR_ox1-silent_, except there was a slight conformational change in the Tyr 367/β and the heterodisulfide product was bound to Ni through its terminal sulfonate. All subsequent MCR structures over the last 25 years have been of the inactive Ni(II) state^9–14^, which has limited our ability to develop structure-based mechanistic proposals.

Attempts at solving the MCR structure in the Ni(I) state have been unsuccessful due to its low redox potential (−0.65 V for the free cofactor)^15^, high oxygen sensitivity^7,16^, and locked-in solvent-inaccessible state precluding in vitro reduction of the Ni(II) form^16^. Even at room temperature, the MCR_red1-silent_ crystal structure shows the “locked in” state containing Ni(II)-thiolate bound HSCoM and HSCoB at 100% occupancy and b-factors similar to those of cryogenic structures^16^. Furthermore, Ni(I)- and Ni(II)-MCR have not been separated by chromatography. Therefore, as described in our methods section, active Ni(I)-MCR is generated by incubating the *M. marburgensis* cell lysate with CO or by harvesting cells in the presence of H_2_^17,18^. Then, to retain the labile Ni(I) state, MCR is kept anaerobic at 4°C in the presence of reductant (Ti-citrate). It is crucial to carefully monitor and validate the fractional composition of Ni(I) to properly interpret structural and mechanistic experiments. Techniques used to monitor the MCR redox state include enzymatic activity, X-ray Absorption and Emission Spectroscopy (XAS, XES), Electron Paramagnetic Resonance (EPR), and UV-Vis spectroscopy. Despite success at generating Ni(I)-crystals^10^, easily identified by their forest green color, for decades our own attempts (using various protocols known to protect other labile systems) have been thwarted, strikingly indicated by immediate conversion of the green color of the crystals to yellow at the beamline. This is because no current crystallography beamline can provide sufficiently anaerobic conditions to keep MCR_red1_ in the Ni(I)-active form.

Therefore, the proposed binding modes and mechanisms of MCR have been heavily influenced by the structures of the inactive Ni(II) protein. However, active MCR appears to lack the Ni-SCoM interaction^3,19^ and is thought to be a dynamic structure, allowing conformational changes to promote catalysis and perhaps downward motion of HSCoB in the substrate channel.^20^ Thus, solving the structure of active Ni(I)-MCR is critical for understanding the mechanism of biological methane formation. Furthermore, because oxidative and radiation damage are of concern with a metalloenzyme this sensitive, it is crucial to rigorously monitor the oxidation state of the crystals during data collection. Here, we describe two anaerobic crystallographic strategies that accomplish these objectives and present the structure of active Ni(I)-MCR using cryogenic synchrotron X-ray diffraction (XRD) and radiation-damage free room temperature serial femtosecond crystallography (SFX) at an X-ray free electron laser (XFEL).

## Results

### Ni(I)-MCR crystal structures by two crystallographic methods validated by spectroscopic and unit cell analyses

A counter diffusion crystallization method was utilized to grow crystals in our anaerobic chamber within a thin quartz capillary, which was sealed with epoxy to provide a transparent vessel for measuring single crystal microspectrophotometry in tandem with room-temperature and cryo-X-ray Diffraction (XRD) (Fig. 1). In addition, we developed anaerobic XFEL serial femtosecond crystallography (SFX) methods with simultaneous X-ray emission spectroscopy (XES) and unit cell measurements^21,22^.

**Figure 1.**
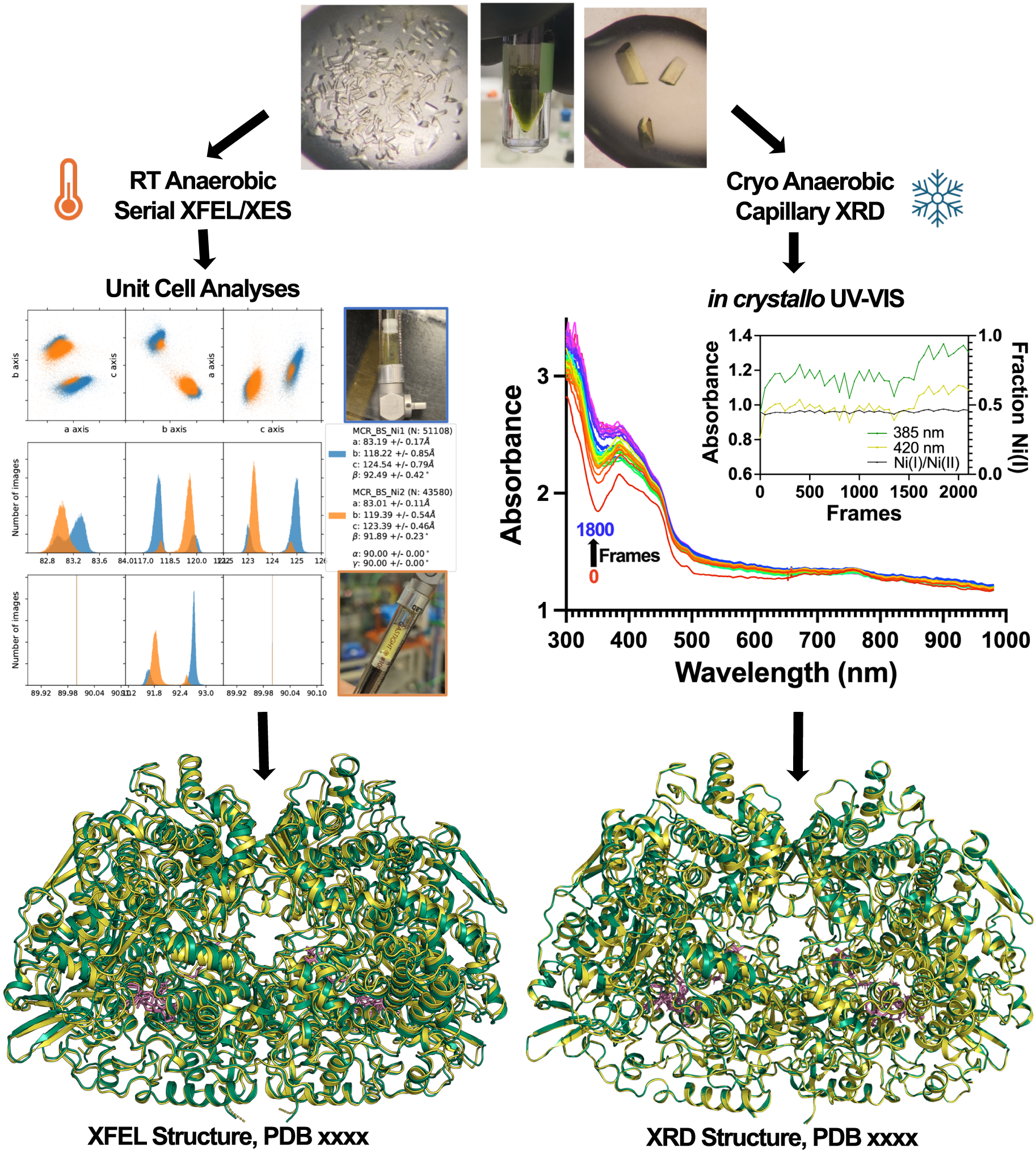
Two methods with two validation tools independently lead to the novel Ni(I)-MCR_red1_. (Top) Green MCR-Ni(I) is crystallized with varying sizes for various methods. (Method, Left) The unit cell distribution of SFX data collection where MCR Ni(I), shown in blue, is found to represent around 77% population of an extended c axis and a shortened b axis. The Ni(I)-MCR sample (Blue border) was removed from the anaerobic chamber and introduced to ambient oxygen. The resulting Ni(II) oxidized sample (Orange border) shifted the unit cell back to the specifications of all previously found Ni(II) crystal structures (Left, Orange peaks). This method results in the structure below where a room temperature SFX crystal structure of MCR obtained from crystals in the Ni(I) unit cell, showing a Ni(I) conformation, represented in green, at an occupancy of 0.54 and a Ni(II) conformation in yellow at an occupancy of 0.46. Cofactor F430, CoM and CoB are represented in purple. (Method, Right) UV-Vis spectra after growth, storage, and data collection of an MCR crystal found to be around 48% Ni(I) (inset). Frames obtained are represented in rainbow where initial frame is red, while frame 2100 is violet. This method results in the cryo-XRD crystal structure of MCR showing two conformations where Ni(I) conformation is represented in green at an occupancy of 0.54, Ni(II) conformation in yellow at an occupancy of 0.46. Cofactor F430, CoM and CoB are represented in purple.

Crystals of MCR 70-75% Ni(I) were grown and housed inside anaerobic 0.1 to 0.3 mm diameter x 20 mm length capillaries^23^. Crystals were analyzed at SSRL Beamline 9-2 by collecting simultaneous XRD and UV-Vis spectral data. The Ni(I) and Ni(II) percentages *in crystallo* stay constant, ranging from 43% to 47% Ni(I) through 2100 frames of data collection, which was representative of all crystals measured. (Fig. 1, Ext. Fig. 2). UV-Vis peaks at 300 and 650 nm also appear during data collection, as seen previously in lysozyme^24^, leading to underlying increases in the amplitude of the 385 and 420 bands. The Ni(I)-MCR was indexed with new unit cell dimensions (a = 83.2 Å, b=118.2 Å, and c=124.5 Å) involving a 1.2 Å shortened b axis and 1.1 Å lengthened c axis (relative to the Ni(II) protein, a = 83.01 Å, b=119.39 Å, and c=123.39 Å) and solved at 1.55 Å resolution.

Room temperature XFEL serial femtosecond crystallography with simultaneous Ni Kα XES was applied to provide a radiation-damage free structure utilizing a serial technique that provides diffraction before destruction^25^. Unlike the SFX data for the yellow Ni(II) crystals grown in the air, those from green Ni(I) crystals (82% Ni(I)) showed two clearly separated clusters of unit cells, one with dimensions of 83.2 Å, 118.2 Å, and 124.5 Å (matching those of the cryogenic Ni(I)-crystal structure) and another matching the Ni(II) structure, indicating crystal packing within MCR depends on the Ni(I)/Ni(II) redox state. A validation tool within the data processing pipeline allows the data to be separately binned based on these unit cell characteristics. The novel unit cell dimension abundance is 77.6%, while the established Ni(II) unit cell is 20.2%. To establish the assignment of the novel unit cell, after Ni(I) data collection, we pulled air into the syringe containing the remaining Ni(I)-crystal slurry and observed a shift in the unit cell from the new form to that associated with the MCR Ni(II) crystal structures. This shift in crystal packing is brought about by a critical ionic bond formation from Lys11/A and Glu123/a (Ext. Fig. 3). Further validation of the redox state of the Ni(I)-crystal slurry was obtained by X-ray emission spectroscopy (Ext. Fig. 4). The Full Width-Half Maximum (FWHM) of the Kα1 line, often used to assess changes in spin and oxidation states^26^, shows a shift of > 0.1 eV between the spectra collected concomitantly with SFX experiments at the XFEL for Ni(I) and Ni(II) crystal samples. This trend is consistent with separate XES measurements on frozen samples of MCR crystal slurries and solutions and on the isolated cofactor F430 at the synchrotron, where an increase by ∼0.3 eV between the Ni(I) and Ni(II) states was observed for deconvoluted samples.

In summary, the cryogenic 1.55 Å XRD and 1.50 Å room temperature SFX crystal structures of Ni(I)-MCR converge (Fig. 1). Using a copy of the model coordinates of the oxidized structure (PDB:7SUC; PDBxxxx, Ext. Fig. 5), an ensemble model was produced including a novel Ni(I) structure conformation plus a Ni(II) conformation to explain the electron densities. Therefore, the maps from both structural methods include two states of MCR, with occupancies of 54% for the Ni(I) and 46% for Ni(II) conformations in which the novel unit cell parameter is found at 77.6% abundance (Ext. Fig 6). It is of note that ensemble refinement used to satisfy the structure factors may not precisely depict the amount of Ni(I) and Ni(II) conformations in the structure. While the two Ni(I) protomers are identical (Ext. Fig. 7), the Ni(I) component of the model is strikingly different from that of Ni(II), which is identical to all past structures and the 1.45 Å Ni(II) SFX structure, generated by oxidation of the Ni(I) crystal slurry. We now describe the large conformational changes that begin at the Ni(I) and F430 cofactor and ripple throughout MCR, extending even to changes in crystal packing reflected in the new unit cell dimensions (These changes are represented in three movies SM. 1-4).

### Nickel Oxidation State Causes Distortion in the NiF430

The first striking feature of the room temperature SFX and cryo XRD structures of the Ni(I) enzyme, is the novel penta-coordinate geometry of the metal ion. A water, located 2.13 Å from the Ni, occupies the lower axial position and replaces the canonical glutamine ligand observed in all Ni(II) structures. CoM relocates with its thiol-sulfur ∼3.9 Å from the metal ion, compared to 2.4 Å in the Ni(II) enzyme. The Ni(I) atom shifts below the F430 plane by ∼1.2 Å compared to the Ni(II) structure, accompanied by significant changes in the F430 ring itself. The N atoms of the corphin in the Ni(II) state are domed relative to Ni(I) (Ext. Fig. 8)^27^. The tetrapyrrole doming leads to a 31° distortion in the C1D-ND-NB-C4B dihedral angle, with a +11° dihedral angle in Ni(I)-MCR compared to the −20° angle for the Ni(II) structure (Fig. 2). In the Ni(I) structure, the dihedral distortion lowers the B ring and alters the positioning of the propionate arms. The results of these changes are also observed in the Ni K-edge XAS (Ext. Fig. 9)^28^. With the one-electron reduction, the EXAFS shows decrease in the Ni-N_F430_ distance by 0.06 Å, replacement of the 2.30 Å Ni-N Gln147/⍺ with a weak axial 2.13 Å water, and loss of the 2.40 Å axial Ni-S (^-^SCoM) bond (Fig. 2). Thus, as shown in SM 1, oxidation of Ni(I) to Ni(II) domes the F430 cofactor, moving the β-lactam upwards and rotating it, which displaces active site Tyr367 of the chain β. These active site movements lead to a seismic cascade through the enzyme.

**Figure 2.**
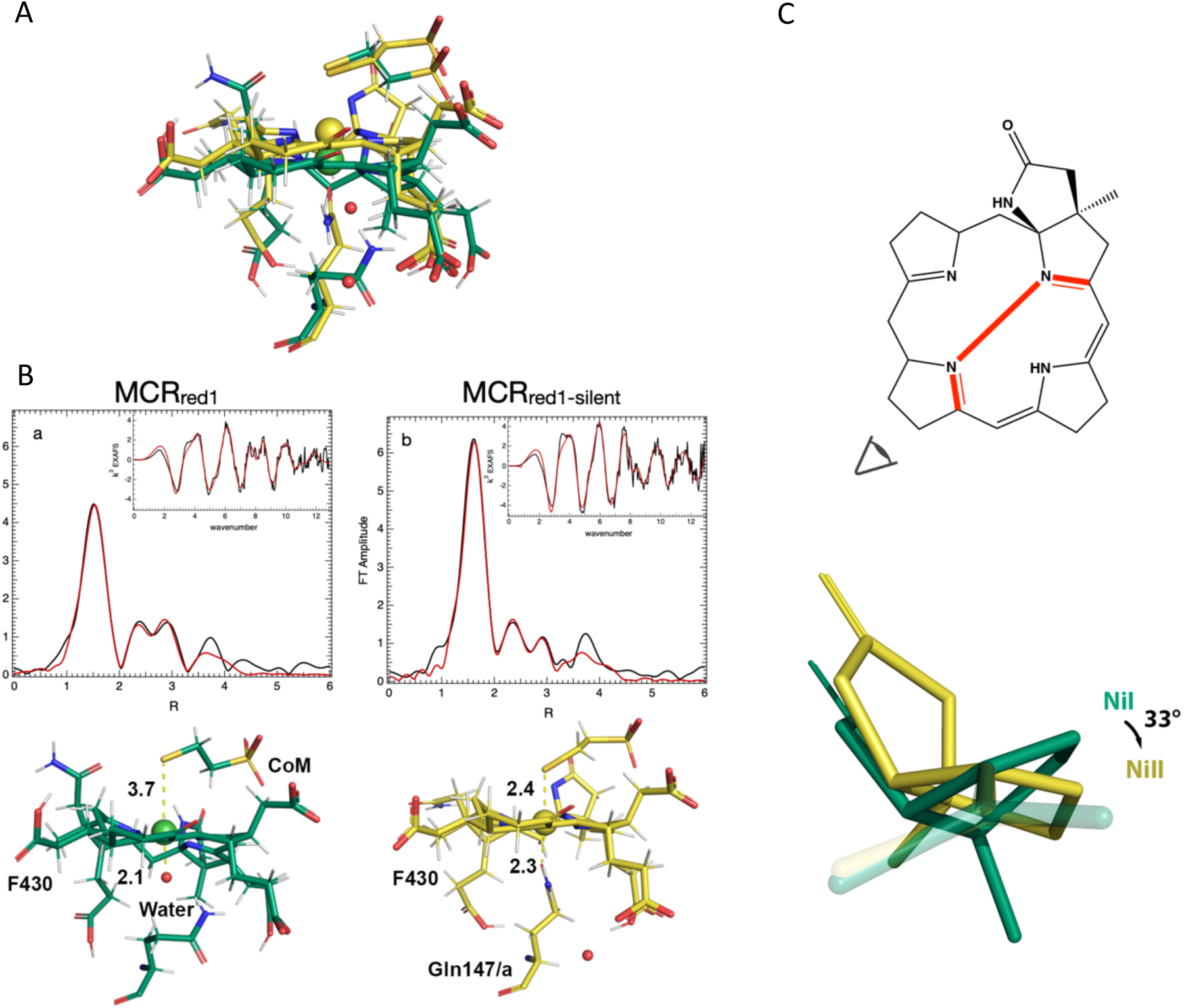
Ni-MCR_red1_ coordination and oxidation state change leads to change in corphin’s β lactam decorated dihedral angle. **A)** Ensemble structure visualizing coordination of both Ni(I) and Ni(II) components of the MCR crystal structure **B)** Ni K-edge EXAFS of MCR_red1_ (a) and MCR_red1-silent_ (b) crystals slurries. Data (black) and fit (red). **C**) Visual of the C1D-ND-NB-C4B dihedral angle from the perspective of the N-N plane illustrating the 33° shift in the dihedral angle. **D**) Refer to Supplemental Movie 1 to compare these changes in coordination environment for the active Ni(I) and inactive Ni(II) proteins.

### Distortion of the Corphin Leads to Clash of the β-lactam Ring and Active Site TYR 367/β

As the β-lactam ring shifts up in response to the doming of the Ni(II)-F430 complex, major clashes occur near the HSCoM binding site (Fig. 3, Movie SM. 2). These include the addition of the 3.0 Å H-bond between HSCoM and the active site Tyr367/β in the Ni(II) structure with a distance of 3.5 Å in Ni(I). Furthermore, in the Ni(I)-enzyme, Arg 130/ɣ exhibits an ionic interaction via the N^+^-^-^oxo with the sulfonate group of HSCoM, strengthened by a hydrogen bond between sulfonate O atom and Arg120/ɣ. In contrast, when oxidized, Ni(II)-MCR locks HSCoM in place with two hydrogen bonding interactions between the sulfonate O atom and the NH_2_ guanidinium moiety of Arg120/ɣ, retaining the other ionic interactions which stabilize HSCoM within the binding site.

**Figure 3.**
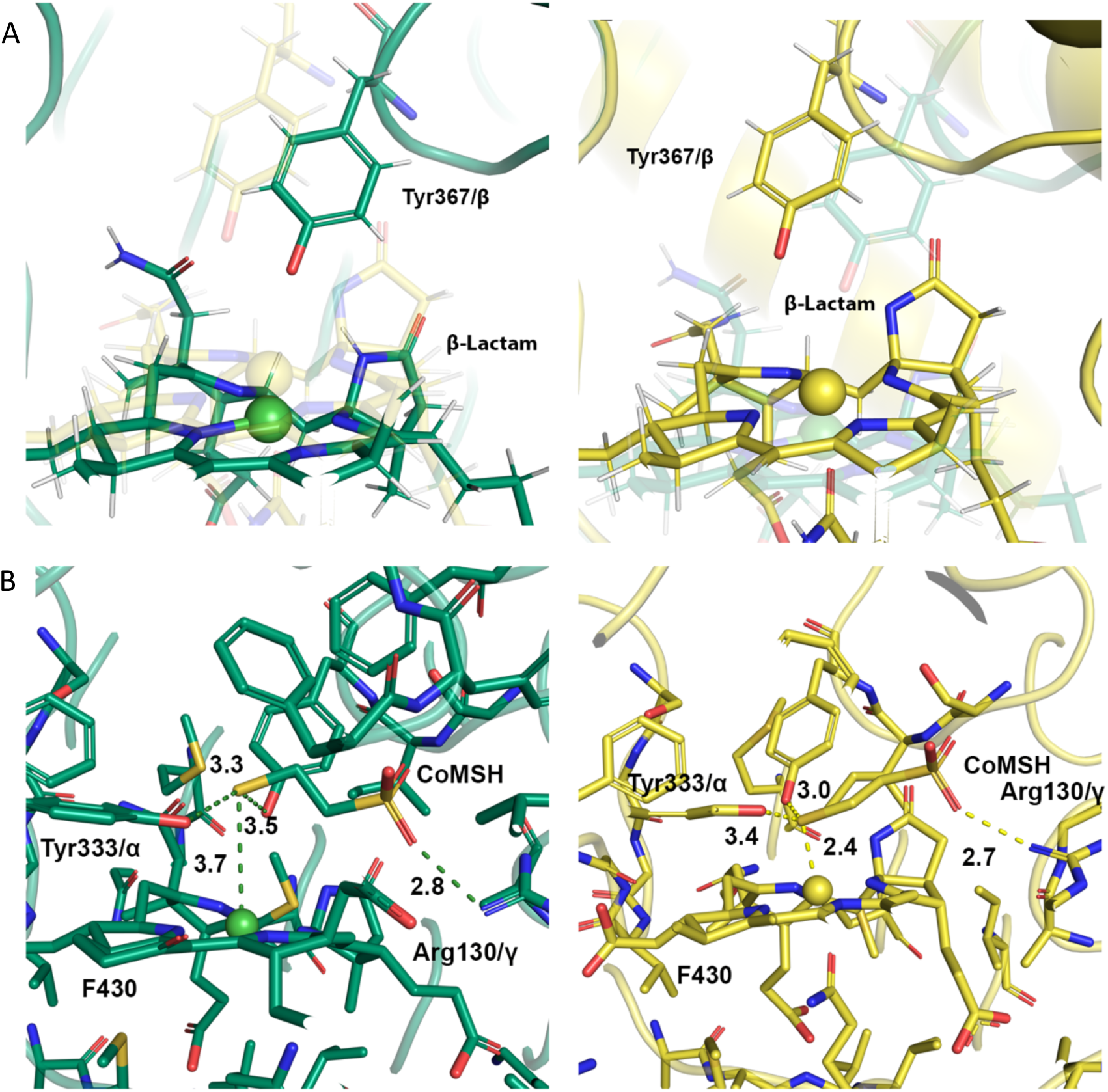
Distortion of the Corphin Leads to Clash of the β-lactam Ring and Active Site TYR 367/β. **A)** Ni(I) to Ni(II) oxidation causes the distortion of the Ni-Cofactor F430 into a doming state moving the β-lactam upwards as well as rotating it with the goal to displace an active site Tyr367 of chain β. **B)** Atomic distances of the Ni(I) model in green and Ni(II) model in yellow. The oxidation positions the thiol of CoM to form a nickel-thiolate bond resulting in a weak 6 coordinate Ni(II) metallocenter. Refer to <u>Supplemental Movie</u> 2 to compare these second sphere changes between the active Ni(I) and inactive Ni(II) proteins.

### TYR 367/β Shifts β, ɣ, and ⍺’ Chains to Close Substrate Tunnel

The movement of Tyr367/β in response to the β-lactam ring initiates relocation of MCR’s entire β chain. His364/β and Tyr367/β are within a flexible loop comprised of residues 364-372, extending into the active site (Fig. 4). In Movie SM. 3, the β-lactam ring-induced movement of Tyr367/β relocates this loop along with His364β, which through an ionic interaction with β-sheet residue Asp113/ɣ, initiates a hinge-like rotation that pulls on the ɣ subunit. Due to the intrachain polar contacts at the β/ɣ subunit interface, these subunits move together, swinging towards the substrate channel at the ⍺, ⍺’, β, and ɣ subunit interfaces. This movement leads to a series of motions that reach the phosphate group of HSCoB, wherein Arg83/ɣ makes ionic contacts (at the ɣ/⍺’ subunit interface) with Glu245/⍺’, within an ⍺-helix (residues 246-257) that also harbors two residues that bind HSCoB, e.g., Lys256/⍺’ and the post translationally modified MHS257/⍺’. Thus, in the Ni(II) form of MCR (Movie SM. 4), the hinge motion pushes this ⍺-helix into the substrate channel, allowing MHS257/⍺’ and Lys256/⍺’ to form ionic interactions with the phosphate group of HSCoB, locking it within the substrate channel (SM. 4). The closing of the substrate channel corroborates that the Ni(II)-MCR structure is indeed a “locked-in” state^16^. Interestingly, this “locked-in” state persists when CoB is absent as evident from the 1.45 Å Ni(II) SFX structure as well as the structure of a 30% CoB-bound MCR by Cederval *et al.*^10^. In the Ni(I) state, the charged clamp at the opening of the substrate tunnel is released, allowing substrate entry and HSCoB to move into position nearer to the CH_3_-S bond of methyl-SCoM.

**Figure 4.**
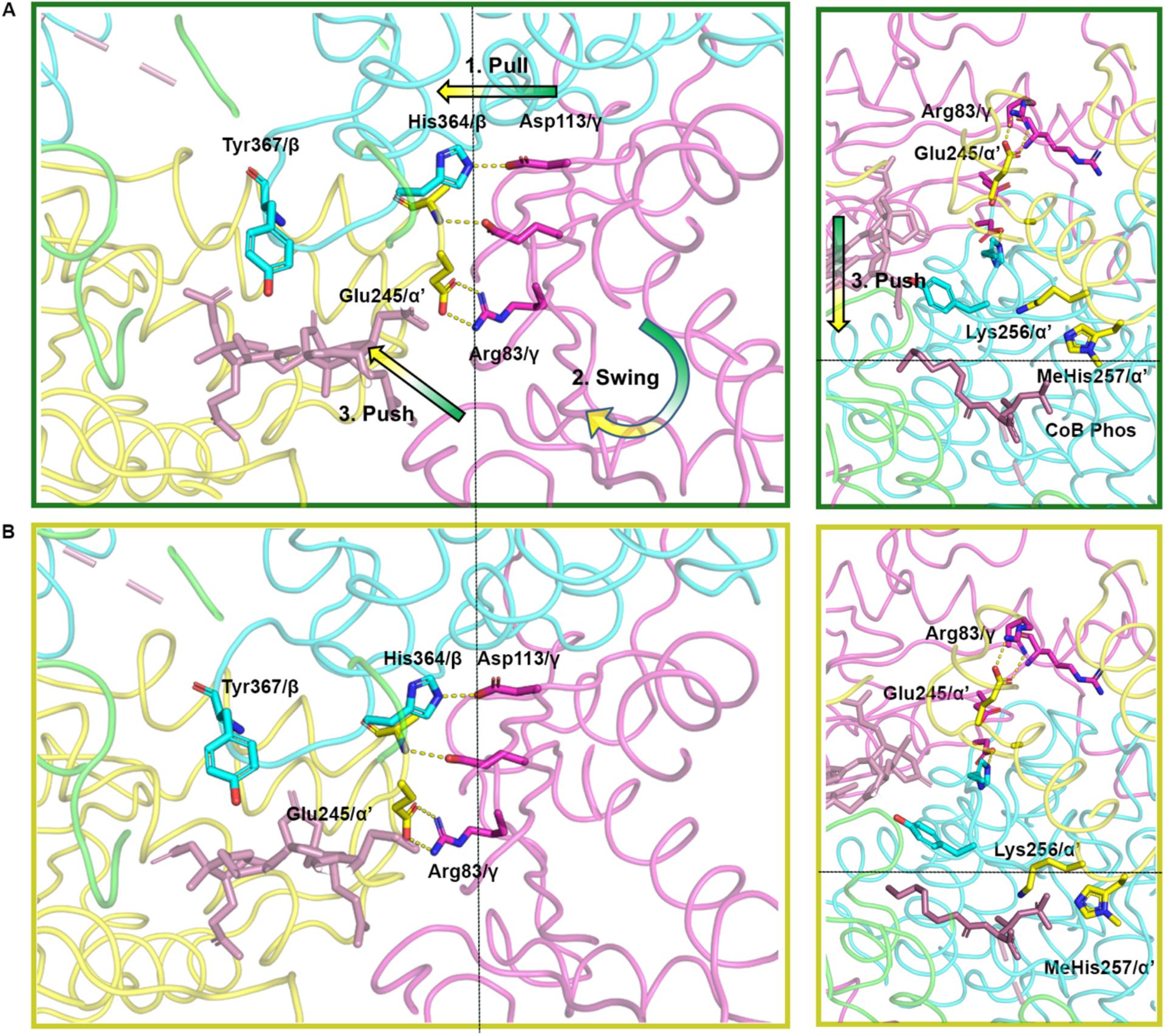
Tyr367/β clash brings global pull-swing-push-close conformational change. MCR conformational changes upon oxidation where the ⍺ subunit is shown in green, the ⍺’ subunit is shown in yellow, the β subunit is shown in blue, and the ɣ subunit is shown in purple. Dotted line used for space reference. Highlighted conformational cause and effect from Ni(I) **(A)** to Ni(II) **(B)** of the Tyr367/β clashing moving the β chain to the left, thus swinging the ɣ chain due to the ionic bond between His364/β and Asp113/ɣ. The swinging of the ɣ chain pushes on the ionic interaction of Arg83/ɣ and Glu245/⍺’ allowing MethylHis257/⍺’ and Lys356⍺’ to bind ionically to the phosphate of CoB. Refer to <u>Supplemental Movie</u> 3 to compare domain movements between the active Ni(I) and inactive Ni(II) proteins (left); and to <u>Supplemental Movie</u> 4 to observe opening and closing the charge clamp (right).

## Discussion

To determine the structure of active MCR, we generated, purified, and crystallized the enzyme in anaerobic chambers maintained at ∼1 ppm O_2_. The major challenge was to retain the Ni(I) crystal and rigorously measure and quantify its redox state during X-ray structure analyses. MCR crystallizes quickly, yields high resolution diffraction patterns, and is amenable to various spectroscopic techniques. Thus, we performed combined XES and SFX in a customized glove box and cryo-XRD on crystals grown and kept in epoxy-sealed quartz capillaries. Quantification of the Ni redox state was accomplished by incorporating the appropriate spectroscopic methods into the structure analysis pipelines. The methods described here for crystallography of MCR are widely applicable to other proteins, especially those with highly sensitive metal centers (e.g., hydrogenase, CODH, acetyl-CoA synthase, nitrogenase). The cryo-XRD and “diffraction before destruction” SFX techniques with tandem validation can be used to further the field of protein crystallography and provide much needed rigorous confirmation of the state of the crystals in structures deposited in the PDB. Thus, for the metalloenzyme and structural biology community, MCR is “a canary in a coal mine”.

There are large differences between the structures of the Ni(I) and Ni(II) proteins. The one-electron reduced Ni(I) protein replaces the lower oxo ligand (from Gln147) with a weak water and lacks the upper thiolate (from CoM). This relaxation and release of the metal-thiolate bond allows for the binding and activation of substrates and removal of substrates, as opposed to all previous Ni(II) crystal structures displaying a “locked-in” inactive state of MCR. This structure provides a clear understanding of the global hinge-like rearrangement that occurs upon one-electron oxidation of penta-coordinate active Ni(I)-MCR. In summary, this oxidation distorts the Ni-Cofactor, causing a 30° upward movement of the β-lactam ring, causing it to clash with Tyr367/β, attached to the β subunit. As Tyr367/β and its accompanying β chain move, the chain tows the ɣ and ⍺’ chains and, through a push-pull mechanism movement involving intra- and inter-subunit contacts, introduce two ionic bonds via MHS257⍺’ and Lys256⍺’ that clamp down on the phosphate group of CoBSH and close the substrate tunnel near the protein surface.

There are important mechanistic implications of how this Ni(I) structure and the conformational changes between the Ni(I)- and Ni(II)-states relate to catalysis. First, the tertiary complex is formed by CH_3_SCoM and CoB binding deep in the substrate binding pocket of the Ni(I) enzyme without the anchoring contacts of the MHS257⍺’ and Lys256/⍺’ (Ext. Fig. 10). The observed release of the Ni-S bond with CoM in the Ni(I) enzyme is a striking contrast with the Ni(II) protein. The presence of Ni-S in the XRD structure of inhibitor-bound Ni(II)-SCoM^9^ was a major rationale for the proposal of a Ni(I)−thioether complex with substrate methyl-SCoM. Patwardhan *et al.* proposed that CH_3_SCoM binds to the active enzyme by a Ni-oxo (sulfonate) instead of a Ni-S (thioether) interaction^19^. Our results support this mechanism by revealing release of the SCoM thiolate to >3.5 Å from the five-coordinate Ni center, in agreement with advanced EPR studies of the Ni(I)-CH_3_SCoM complex^19,20^. Thus, the Ni(I)-thiolate forms a weaker bond than Ni(II) (and the Ni(I)-thioether of methyl-SCoM would be even weaker), leading to protonation of SCoM (pK_a_ = 9.4)^19^ and formation of another H-bond with Tyr367/β. These events are triggered by the cascade initiated by movement of the β-lactam ring of F430. The proposed Ni(I)-OSO_2_ binding mode is additionally supported by EPR studies using ^33^S-substituted H^33^SCoM and by near-infrared (near-IR), XAS, and EPR spectra, which ruled out a Ni-sulfur interaction in the complex of Ni(I)-MCR with HSCoM.^19,20^

The observed >3.5 Å Ni(I)-SCoM distance supports the proposal of a long-range electron transfer in generation of a methyl radical that will abstract a H atom from HSCoB.^19^ We speculate that CH_3_SCoM has two competing binding modes, entailing a dynamic Arg120 that flips in and out of its hydrogen bonding interaction (with SCoM sulfonate) and the thioether of CH_3_SCoM acts as a weaker H-bond acceptor and weakly coordinates with the Tyr333/β and the dynamic Tyr367/β, allowing for a less constrained CH_3_SCoM.

In the first catalytic step in the MCR mechanism (Ext Fig. 10), electron transfer from Ni(I) initiates cleavage of the S-CH_3_ bond of CH_3_SCoM to generate Ni(II) and a methyl radical intermediate. Assuming the observed Ni(II) state mimics this Ni(II)/methyl radical state, we speculate that reaction involves the global movement described above to close the substrate channel via MHS257⍺’ and Lys256/⍺’. These interactions may be dependent on HSCoB binding as seen in our Ni(II) oxidized SFX structure (Ext Fig 5) and in Cederval *et al*^10^. Homolytic bond cleavage then creates the transition state of Ni-S-CH_3_-HSCoB. Upon hydrogen atom abstraction, MCR undergoes the redox induced butterfly effect causing HSCoB to shift upward and create the canonical ionic interactions with MHS257/⍺’ and Lys256/⍺’ as well as guided movement of the Tyr367/β hydrogen bonding interactions to SCoM. The CoBSᐧ radical would then react with the HSCoM thiol creating the disulfide anion radical. Lastly, the disulfide anion radical reduces the Ni(II) via long range electron transfer, to generate CoBSSCoM, which is released to complete turnover. Note that these reactions occur in reverse for the reverse methanogenesis reaction.

These dynamics spring from an unexpected structural change in F430, which is a unique cofactor specific for methanogenic and methanotrophic archaea that use MCR for methane formation and oxidation, respectively^7^. It is an evolutionary cousin of heme and B_12_, but its Ni(II)/Ni(I) couple, between −600 and −700mV vs. NHE, is the lowest of any cofactor, as well as being the only naturally occurring Ni-tetrapyrrole known^15^. F430 includes several propionate arms along with two unique cyclic structures, a β-lactam ring beside tetrapyrrole ring B and a keto-containing carbocyclic ring. Both the propionates and the β-lactam rearrange between the Ni(I) and Ni(II) structures. Additional functionalization of the tetrapyrrole and structural constraints imparted by the oxidation state of the bound metal are responsible for distortions like ruffling, saddling, waving, or doming in other well-known tetrapyrrole cofactors, like heme, tuning the electronic structure and reactivity of the complex^29,30^. These tetrapyrrole nuances fine tune the chemistry alongside the large conformational changes related to the enzyme mechanism established in this work.

Our results utilizing MCR, the canary in the coal mine, illustrate a strategy for generating, maintaining, and validating labile states of macromolecular crystals during data collection. In this study, we used in-line UV-visible microspectrophotometers for *in-crystallo* monitoring of the Ni redox state for the synchrotron experiments as well as XES for the SFX room temperature experiments. The validation of oxidation state *in crystallo*, which is crucial for mechanistic interpretation, is rarely employed for structures deposited into the PDB. The findings presented here underscore the importance of such validation, and as such we call on the structural biology community to provide rigorous validation of the (oxidation) state of the crystal upon structure deposition into the PDB. To rigorously validate the Ni(I) structure of MCR, we combined room-temperature “diffraction before destruction” SFX in addition to cryo XRD. This Ni(I) structure provides the first snapshot of the active state of MCR, revealing and tracking the large conformational changes relative to the locked-in Ni(II) state. The active enzyme has an open upper axial coordination site poised for binding and reaction with substrates. This work poises MCR research for important next steps, e.g., trapping substrates and analogs in the binding pocket of the active enzyme and determining the structures of Ni(I) variants using developments in site-directed mutagenesis^13^. A pressing question that remains is what (e.g., dynamic thermal networks^31^) provides the spark to initiate catalysis.

## Materials and Methods

### Organism and Materials

*Methanothermobacter marburgensis* was obtained from the Oregon Collection of Methanogens (Portland, OR) catalog as OCM82 and cultured on H_2_/CO_2_ (80/20%) at 65 °C as described^19^. All buffers, media ingredients, and other reagents were acquired from Sigma. N_2_ (99.98%), CO (99.99%), H_2_/CO_2_ (80%/20%), and Ultra High Purity (UHP) H_2_ (99.999%) gases were obtained from Cryogenic Gases (Grand Rapids, MI). A stock solution of 83 mM Ti(III) citrate was prepared^19^, and substrates, methyl-SCoM^32^ and CoBSH^19^ were synthesized and purified, as described.

### MCR Purification and Crystallization

Ni(I)-MCR_red1_ (the MCR-1 enzyme) from *M. marburgensis* (catalog OCM82) was activated in the cell lysate with CO, purified and handled in an anaerobic chamber (Vacuum Atmospheres, Inc., Hawthorne, CA) under a N_2_ atmosphere containing <0.2 ppm of O_2_, as previously described^19^. Before crystallization, Ni(I)-MCR_red1_ was prepared with five rounds of buffer exchange to remove HSCoM and salt from the storage buffer. UV-Vis was used to quantify the Ni(I) percentage using extinction coefficients of 22.0 and 12.7 mM^-1^ cm^-1^ at 420 and 385 nm, respectively, with a previously described multiwavelength calculation^10,19,33^.

Crystallization and preparation of MCR for XRD and SFX measurements were performed by different protocols. Batch crystallization of MCR for SFX was carried out by mixing 5 μL of 1.5 mM Ni(I)-MCR_red1_ (400 mg/ml) and 45 μL of crystallization buffer (100mM HEPES pH 7.5, 150mM Magnesium Acetate, 18% PEG 400) with a seed stock concentration previously determined to result in small 20-80 μm sized crystals. Crystals formed in 1-2 days. For XRD, capillary crystallization by counter diffusion was utilized where 0.1 – 0.3 mm diameter x 20mm length capillaries were filled with a batch prep crystallization mixture described above^23^. Capillaries were then placed in crystal buffer with an agarose plug and left to crystalize overnight. After growth, capillaries were placed in a high PEG 400 cryo-protectant buffer (100 mM HEPES pH 7.5, 150 mM Magnesium Acetate, 25% PEG 400) for 36 hours. Epoxy was used to seal the capillaries on each side and to adhere the capillary to a MiTeGen B1 mount. The mounted capillaries were taken out of the anaerobic chamber and cryofrozen by quenching and then stored in liquid nitrogen until shipment to SSRL.

### Serial Femtosecond X-ray Diffraction (SFX) and X-ray Emission Spectroscopy (XES) data collection at LCLS

SFX diffraction data were collected at the Macromolecular Femtosecond Crystallography (MFX) instrument of LCLS, (SLAC National Accelerator Lab, Menlo Park, CA)^34^ at 300 K on a RAYONIX MX340-HS CCD detector, using the previously established Drop-On-Tape (DOT)^22^ approach. The crystal slurries were loaded in gas tight Hamilton syringes which were placed on a sample rocker inside an anaerobic chamber which maintained an oxygen level of <5ppm and a temperature of 15C throughout the experiment. The DOT setup itself was in a separate He filled enclosure and maintained at 400 ppm O_2_ and 26C. The capillary line connecting the Hamilton syringe to the sample reservoir inside the DOT setup was enclosed by a cooling sleeve and kept at 12C. A sample flow rate of 6 μl/min was utilized resulting in droplets of 3.3 nl. The belt speed was 300 mm/s, hence the travel time of droplets from dispension to the X-ray interaction point was 0.8 s. The X-ray beam photon energy was 9.5 keV with a pulse energy of 2-3 mJ, a pulse length of 35 fs and a beam size on the sample of 2.5 μm x 2.5 μm (Full Width Half Max, FWHM). Data collection statistics are available in SI Table 1.

X-ray emission data were collected in tandem with diffraction data using a multicrystal wavelength-dispersive hard X-ray spectrometer based on the von Hamos geometry^35^. Due to the change of polarization direction of the hard X-ray undulator of LCLS our previously used setup^22,35^ was modified to place the analyzer crystals above the X-ray interaction point and the position sensitive detector at 90 degrees from the beam direction in the horizontal plane. An array of three Si(620) analyzer crystals was placed 250 mm above the interaction point with the center of the crystals at 74.80 degrees with respect to the interaction point, collecting both Ni Kɑ lines on an ePix 100 detector with its center located 136 mm side wise of the X-ray interaction point. To calibrate the spectrometer geometry, spectra from [Ni(H2O)6]^2+^ and a Ni metal foil were collected and compared to a synchrotron reference. The XES data was also pedestal corrected to account for differences in noise of the detector pixels. For the background subtraction, a slice on either side of the region of interest (ROI) was selected. A row-by-row first order polynomial fitting scheme was utilized for the initial two-dimensional background subtraction. To then bring the baseline of the spectra to zero, a one-dimensional background subtraction (also utilizing a first order polynomial fitting scheme) was also deployed. The spectra were smoothed using a Savitzky-Golay filter with a window length of 9 and polynomial order of 3 and subsequently were area normalized.

### SFX Data Processing

The collected dataset was reduced and processed using *cctbx.xfel* and DIALS^36,37^. We performed joint refinement of the crystal models against the detector position for each batch to account for small time-dependent variations in detector position^37^. The two samples were found to be a mixture of two isoforms with the dominant isoform swapped between them. Prior to scaling, for each sample, a unit cell clustering analysis was performed to separate the two isoforms (SI Fig. 1). We used the clustering algorithm implemented in *cctbx.xfel.merge* to select only the cluster of interest during scaling/merging with a mahalonobis parameter set to 4.0. The results from the clustering as well as the ε parameters used are also shown in SI Fig 1. Data was scaled and merged to 1.5 Å (Ni(I)-MCR) and 1.45 Å (Ni(II)-MCR), respectively, for the two samples based on previously established resolution cutoff criteria (∼10× multiplicity, where the values of I/σ(I) do not uniformly decrease any more, and where cc1/2 values stop decreasing monotonically, indicating no useful information is contained in resolution shells beyond that point), using cctbx.xfel.merge with errors determined by the mm24 method^38^ as available on the cctbx GitHub page [https://github.com/cctbx/cctbx_project] which is a modification of the ev11 error model^22,39,40^. This procedure uses concepts from robust statistics to more reliably calibrate intensity uncertainty estimates with non-normal error and outlier observations. Absorption correction from the kapton tape is not performed. This absorption is a relatively small error compared to partiality^41^ and more accurate correction procedures would be needed to account for this error. Data statistics are available in SI Table 1. Structure determination was done using Phenix^42^ starting with molecular replacement using PDB ID: 7SUC^16^ as the reference model^43^. For all subsequent refinements with Phenix, we turned off automatic linking within the chain, as well as NCS restraints, and instead defined the interactions between the Ni-OE1(Q147) and Ni-S1(CoM) by supplying custom coordination restraints as parameter (phil) files. We used Coot^44^ for model building with multiple iterations of refinement using *phenix.refine*^45^. All figures for this paper were generated using Pymol^46,47^.

### X-ray Diffraction and UV-Vis data collection at SSRL

X-ray diffraction data were collected at SSRL beamline 9-2 equipped with a PILATUS 6M PAD (Dectris, Switzerland) at a wavelength of 0.98 Å. Data was collected at 100K, 75 x 100 um beam size, 579 mA beam flux, and 0.4 s exposure time. The data was processed with xia2 and DIALS^36,48^. Molecular replacement was carried out with Phaser-MR in the Phenix suite and refined with phenix.refine^49,50^ by iterative modelling using Coot^44^. UV-Visible Absorption micro-Spectrophotometry (UV-Vis AS) at SSRL BL9-2 was employed to characterize the starting MCR oxidation state and track the effect of radiation damage on the redox center during X-ray data collection. The Microspec was focused to a 40 µm diameter spot to the crystal and the angle that produced the clearest absorbance results was determined to be 155 degrees. X-ray data collection was interleaved with absorption spectra at this angle every 90 frames/9 degrees of rotation.

### Ensemble Refinements

Molecular replacement was employed using PHASER-MR via the Phenix suite^50^ to solve phases using 7SUC phase information for both SFX structures of Ni(I)-MCR and air oxidized Ni(II)-MCR as well as the Ni(I) MCR cryogenic structure. The Ni(II)-MCR SFX structure was refined and solved first. The Ni(II)-MCR coordinates were positioned into the Ni(I)-MCR SFX electron density and a new conformation of the protein was created to form an ensemble. The subsequent Fo-Fc maps were used to ascertain the Ni(I) conformation of the Ni(I)-MCR SFX ensemble structure. Occupancies of the conformations were grouped, and occupancy refinement was used to adjust the occupancy of the two conformations. The occupancy obtained this way and the percentage of Ni(I) based on ex-situ UV-Vis measurements on the same sample preparation support each other independently. The cryogenic Ni(I)-MCR structure was then modelled based on the SFX Ni(I)-MCR ensemble structure and the occupancy cross checked with the in-line microspec measurements on the same crystal.

### X-ray Absorption Spectroscopy

MCR crystals were prepared as described above. Ni(I)-MCR_red1_ crystals were loaded into solution XAS cuvettes and sealed with 25 μm Kapton tape under anoxic conditions using a N_2_ glove box, following which, the cuvettes were flash frozen and shipped to SSRL under LN_2_. XAS spectra were measured on the cuvettes within 24 hours of shipment. Ni(II)-MCR_red1-silent_ crystal slurry was shipped separately at room temperature, and syringed into XAS cuvettes, which were then kept under LN_2_ during handling. Ni K-edge XAS were collected at the Stanford Synchrotron Radiation Lightsource (SSRL) on beamline 9-3, a 2T 20-pole wiggler side station. Fluorescence data were collected using a Canberra 100-element monolith Ge detector. Energy selection was provided by a Si(220) double-crystal monochromator oriented to ϕ = 0, and harmonic rejection was provided by a spherically bent, Rh-coated mirror. The beamline was optimized to the 10 KeV cutoff to prevent higher-order harmonics from contaminating the signal. Unwanted signal from photon scattering was reduced using a Co filter and Soller slits. During data collection, samples were maintained at 10 K using an Oxford Instruments CF 1208 liquid He cryostat. Energy calibration for each scan was obtained from the simultaneous collection of scans on a Ni foil which was placed between the second and third ion chambers. Spectra were collected at multiple spots per sample to minimize beam-induced damage, although no beam damage was observed across multiple scans on the same sample spot.

Due to the dilute nature of the samples, and because a large background signal from diffraction (both from ice and the protein crystal matrix) was observed in some of the channels, the fluorescence data were analysed channel-by-channel (each channel representing a single detector element of the 100-element Ge detector) using Larch^51^ to identify and exclude the compromised channels. Channels that were found to be satisfactory were then summed for each scan and imported to Athena^52^ for calibration and normalization. Spectra of the reference foil were calibrated to 8331.6 eV, and the energy shift was then applied to the corresponding sample spectrum. The data were normalized by applying a linear background function to the pre-edge region, and a third order polynomial to the post-edge region. To extract the EXAFS signal, the data were transferred to Pyspline^53^ where the EXAFS spectra were extracted by setting the value of E_0_ to 8345 eV and fitting a 4-region spline to the data with polynomial orders 2, 3, 3, and 3. Least-squares fitting of the EXAFS data was done in Artemis^52^. Feff8^54^ was used to generate theoretical EXAFS phase and amplitude parameters of scattering paths within a 5 Å radius of the Ni absorber within their respective crystal structures. Four parameters were evaluated for each backscattering path in course of fitting: the interatomic distance R, the interatomic disorder σ^2^, the shift in the assigned value of E_0_, and the amplitude reduction factor S_0_^2^. Of these four parameters, R and σ^2^ were allowed to float for each interaction, while the shift in E_0_ was allowed to float but fixed to a common value among all paths of a given fit. S_0_^2^ was fixed at 1.0. Peak deconvolution was done in Larch v9.45 using pseudo-Voigt functions to model the Ni K-pre-edge feature.

### X-ray Emission Spectra collected at SSRL

Ni K⍺_1,2_ X-ray emission spectra of MCR crystal slurries as well as of MCR solution samples and solutions of the isolated F430 cofactor were collected at SSRL beamline 15-2 at 10K under standard ring operating conditions. The beam was focused using KB mirrors to 740 µm x 190 µm, and energy selection was provided by a liquid N_2_ cooled Si(311) double crystal monochromator. Spectra were collected using a 7-crystal Johann X-ray spectrometer^55^. The incident X-ray energy was set to ∼8600 eV, and the Ni K⍺_1,2_ region emission was scanned from 7485 eV to 7450 eV. Data were processed and analyzed in MATLAB or jupyter notebooks, where a linear background was subtracted from the spectra, and normalization achieved by scaling to a unit integrated intensity. For the crystal slurry samples in order to account for the 31% of MCR_red1-silent_ present in the MCR_red1_ sample, a subtracted spectrum was made, which was then renormalized to unit area for comparison. This corrected spectrum is denoted by an *. The Ni(I)-MCR_red1_ and Ni(II)-MCR_red1-silent_ samples were both prepared from the same preparation, starting out as MCR_red1_, and the MC_Rred1-silent_ sample was made by allowing an aliquot of sample to oxidize in air until fully converted, and confirmed by UV-Vis spectroscopy, allowing for the subtraction. The FWHM of the data were calculated numerically from the K⍺_1_ line, rather than from peak fitting, due to poor agreement between the data and common line shapes. However, the numerically calculated values, and those obtained from least-squares fitting with a pseudo-Voigt are comparable (Ext. Fig. 4).

## Author Contributions

C.J.O., D.G.V. prepared samples

M.J.A., R.S., H.M., M.D. performed XAS and XES experiments at SSRL.

C.J.O., D. M.-R., A.E.C, S.W.R. performed XRD experiments at SSRL.

C.J.O., A.B., K.C., H.M., M.D., D.M., D.W.P., N.W.M., N.K.S., A.S.B., J.F.K. processed and analyzed SFX data.

P.S.S., K.C., H.M., V.T., H.S., J.F.K, developed, tested and ran the SFX sample delivery system. P.S.S., H.M., K.C., L.B.G., R.A.-M., U.B., V.K.Y., J.Y., J.F.K. developed and tested the XES setup at LCLS.

V.T., H.S., D.J.R, L.B.G, R.A.-M. operated the MFX instrument.

C.J.O., P.S.S., A.B., K.C., H.M., V.T., S.K., V.K.Y., J.Y., J.F.K., S.W.R. performed the SFX experiment at LCLS.

C.J.O., S.W.R. wrote and edited the manuscript.

J.F.K., R.S., and M.J.A. edited the manuscript.

## Supporting information

Ext. Fig.

## Acknowledgements

We want to thank Anjali Patwardhan for the protocols and discussions contributing to this work. We thank Isabel Bogacz and Corey Kaminsky for help during some of the synchrotron XES measurements. The work was supported by the U.S. Department of Energy, Office of Science (OS), Office of Basic Energy Sciences (BES), Chemical Sciences, Geosciences, and Biosciences Division, Physical Biosciences Program through DE-FG02-08ER15931 (S.W.R.), FWP 100593 (R.S.) and Contract No. DE-AC02-05CH11231 (J.Y., V.K.Y., J.F.K.) and by the National Institutes of Health (NIH) Grants GM149528 (V.K.Y.), GM110501 (J.Y.), GM126289 (J.F.K.), GM117126 (N.K.S.), GM151988 (N.K.S.). C.J.O was supported in part *by* the U.S. Department of Energy, Office of Science, Office of Workforce Development for Teachers and Scientists, Office of Science Graduate Student Research (SCGSR) program. The SCGSR program is administered by the Oak Ridge Institute for Science and Education for the DOE under contract number DE-SC0014664.

Research was carried out at the Linac Coherent Light Source (LCLS), SLAC National Accelerator Laboratory (proposal no. L10100 and L10297), supported by the DOE Office of Science, OBES under Contract No. DE-AC02-76SF00515. SFX data processing was performed in part at the National Energy Research Scientific Computing Center, supported by the DOE Office of Science, Contract No. DEAC02-05CH11231.

The Rayonix detector used at LCLS was supported by the NIH grant S10 OD023453. Data processing was supported by the US DIALS National Resource (R24GM154040). Experiments at the LCLS were supported by the NIH grant P41GM139687. Use of the Stanford Synchrotron Radiation Lightsource, SLAC National Accelerator Laboratory, is supported by the U.S. Department of Energy, Office of Science, Office of Basic Energy Sciences under Contract No. DE-AC02-76SF00515. The SSRL Structural Molecular Biology Program is supported by the DOE Office of Biological and Environmental Research, and by the National Institutes of Health, National Institute of General Medical Sciences (P30GM133894). The contents of this publication are solely the responsibility of the authors and do not necessarily represent the official views of NIGMS or NIH.

## Notes

### Competing Interest Statement

The authors have declared no competing interest.

