## Supplementary material for "Structure of active methyl-CoM reductase, Earth’s main methane producer": SI Table I, SI Fig. 1

*Supporting Info*

*Tables*

| ***Data set*** | **Ni(I) XFEL** | **Ni(II) XFEL** | **Ni(I) Cryo** |
|---|---|---|---|
| ***Data Collection*** | | | |
| *Resolution range (Å)* | 20.49-1.5 | 20.44-1.45 | 41.29-1.55 |
| *Resolution upper bin (Å)* | 1.53-1.50 | 1.48-1.45 | 1.61-1.55 |
| *Wavelength (Å)* | 1.262 | 1.262 | 0.979 |
| *Space group* | P1211 | P1211 | P1211 |
| *Unit cell parameters (Å)* | a=83.32 | a=83.04 | a=82.78 |
| | b=117.89 | b=119.61 | b=117.49 |
| | c=125.03 | c=123.29 | c=123.99 |
| *Lattices merged* | 33937 | 30804 | 2070 |
| *Unique reflections* | 382644 | 423896 | 333953 |
| *(upper bin)* | 17958 | 20950 | 31743 |
| *Completeness* | 99.57 | 99.92 | 94.07 |
| *(upper bin)* | 93.7 | 99.29 | 61.22 |
| *CC1/2* | 0.992 | 0.992 | 0.998 |
| *(upper bin)* | 0.199 | 0.2843 | 0.48 |
| *$I/\sigma(I)$* | 5.26 | 5.95 | 8.97 |
| *(upper bin)* | 0.43 | 0.64 | 0.28 |
| *Multiplicity* | 204.07 | 224.27 | 4.02 |
| *(upper bin)* | 4.59 | 7.54 | 3.28 |
| *Wilson B-factor* | 22.22 | 19.65 | 24 |
| ***Refinement*** | | | |
| *Resolution range (Å)* | 20.49-1.50 | 20.44-1.45 | 41.29-1.55 |
| *Resolution upper bin (Å)* | 1.53-1.50 | 1.48-1.45 | 1.61-1.55 |
| *R-factor* | 0.1305 | 0.128 | 0.1462 |
| *R-free* | 0.1709 | 0.148 | 0.1978 |
| *Number of atoms* | 77281 | 41170 | |
| *Number non-hydrogen atoms* | 39922 | 21527 | 40652 |
| *Ligands* | 13 | 11 | 12 |
| *Waters* | 1329 | 205 | 1969 |
| *Protein residues* | 2476 | 2476 | 2476 |
| *RMS (bonds)* | 0.008 | 0.009 | 0.01 |

| | | | |
|---|---|---|---|
| *RMS (angles)* | 0.959 | 1.036 | 1.09 |
| *Ramachandran favored* | 96.4 | 97.2 | 94.77 |
| *Ramachandran outliers* | 0.28 | 0.15 | 0.41 |
| *Clashscore* | 2.88 | 1.71 | 8.95 |
| *Average B-factor* | 31.46 | 31 | 29.58 |

*Methods*

*X-ray absorption spectroscopy of MCR crystals*

Ni K-edge XAS spectra were measured on slurries of MCR crystal in the $MCR_{red1-silent}$ and $MCR_{red1}$ states. UV-Vis spectra taken on the species crystalized exclusively for XAS analysis revealed 14.6% Ni(II) present in $MCR_{red1}$. To correct for this, the spectra of the pure $MCR_{red1}$ was obtained by subtracting out the $MCR_{red1-silent}$ signal weighted by 0.146. These spectra are presented in **Ext Fig 3**. Each spectrum exhibits a pre-edge feature arising from the electric dipole forbidden, quadrupole allowed 1s → 3d transition, occurring at ~8331.4 eV and ~8331.9 eV in the $MCR_{red1}$ and $MCR_{red1-silent}$ species, respectively. $MCR_{red1}$ also had an additional feature composed of two peaks on the rising edge at 8333.4 and 8334.9 eV, obtained from peak deconvolution **Ext Fig 4**. This type of feature is most intense in complexes with four coordinate $D_{4h}$ symmetry where substantial metal to ligand back-bonding is present and has been previously observed in solution-phase $MCR_{red1}$[1].The position of the pre-edge and rising edge transitions is in excellent agreement with prior measurements on solution-phase protein[1]. When oxidized to $MCR_{red1-silent}$, the pre-edge position increases in energy by ~0.5 eV, in agreement with the solution-phase data [1] indicative of an increase in ligand field strength accompanying an increase in coordination number. The amplitude of the pre-edge feature in $MCR_{red1-silent}$ is greater than in $MCR_{red1}$, reflecting the differences in coordination, and is consistent with the earlier solution-phase data. In the white line region (8340-8360 eV), major differences are also present across the sample series reflecting the differences in geometry.

EXAFS were collected to K = 12.5 $Å^{-1}$ on all samples and were fit using the co-collected crystallographic structures as input models. The EXAFS and corresponding Fourier-transform data are presented in **Ext Fig 3**, and details of the best-fits are also presented in **Ext Fig 3**. The $MCR_{red1}$ is best fit as five-coordinate, with four N from the corphin ring at **2.03 Å** and one O at **2.16 Å**, while the $MCR_{red1\text{-}silent}$ is best modelled as a six coordinate species, with the $Ni\text{-}N_{F430}$ elongated to **2.08 Å**, the O from the axial Gln at **2.18 Å** and the S of the bound SCoM at **2.43 Å**. To account for the approximately 15% Ni(II) present in $MCR_{red1}$, a S path was also fit, weighted by a coordination number of 0.15. These distances are in excellent agreement with prior solution phase data[1], as well as the crystallographic data. As with the solution structure data, modelling the $Ni\text{-}N_{F430}$ contribution of $MCR_{red1}$ requires a relatively large value of $\sigma^2$ factor, indicative of heterogeneity in the Ni-N bond distances, as implicated by the distortions of the corphin ring exhibited by the crystal structure, and analyzed with PorphyStruct[2] as shown in Fig. 2 in the main text.

*Figures*

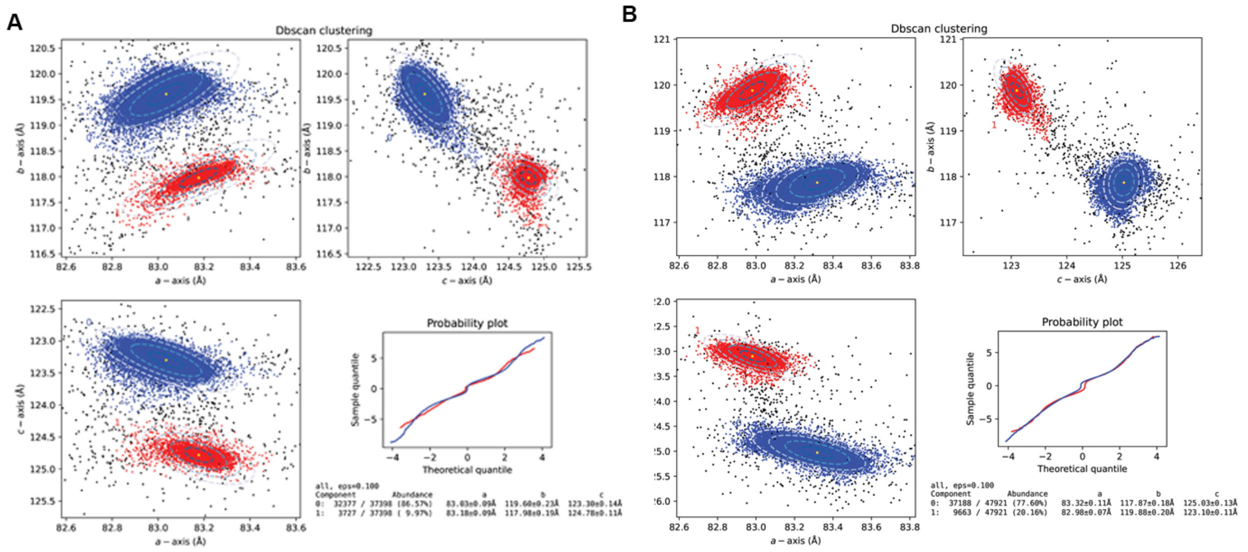

Supplemental Figure 1. Unit cell clustering analysis performed on the MCR data as implemented in *cctbx.xfel.merge* program for sample (A) Ni(I)-MCR and after oxidation (B) Ni(II)-MCR. The clustering step helps separate out the isoform of interest in the respective samples prior to scaling/merging. It is noteworthy that the dominant isoforms are swapped between the two samples.

1 Sarangi, R., Dey, M. & Ragsdale, S. W. - Geometric and Electronic Structures of the NiI and Methyl−NiIII Intermediates of Methyl-Coenzyme M Reductase. **- 48**, - 3156 (2009).

2 Krumsieck, J. & Bröring, M. PorphyStruct: A Digital Tool for the Quantitative Assignment of Non-Planar Distortion Modes in Four-Membered Porphyrinoids. *Chemistry – A European Journal* **27**, 11580-11588 (2021). https://doi.org/https://doi.org/10.1002/chem.202101243
