## Supplementary material for "Structure of active methyl-CoM reductase, Earth’s main methane producer": Ext. Fig.

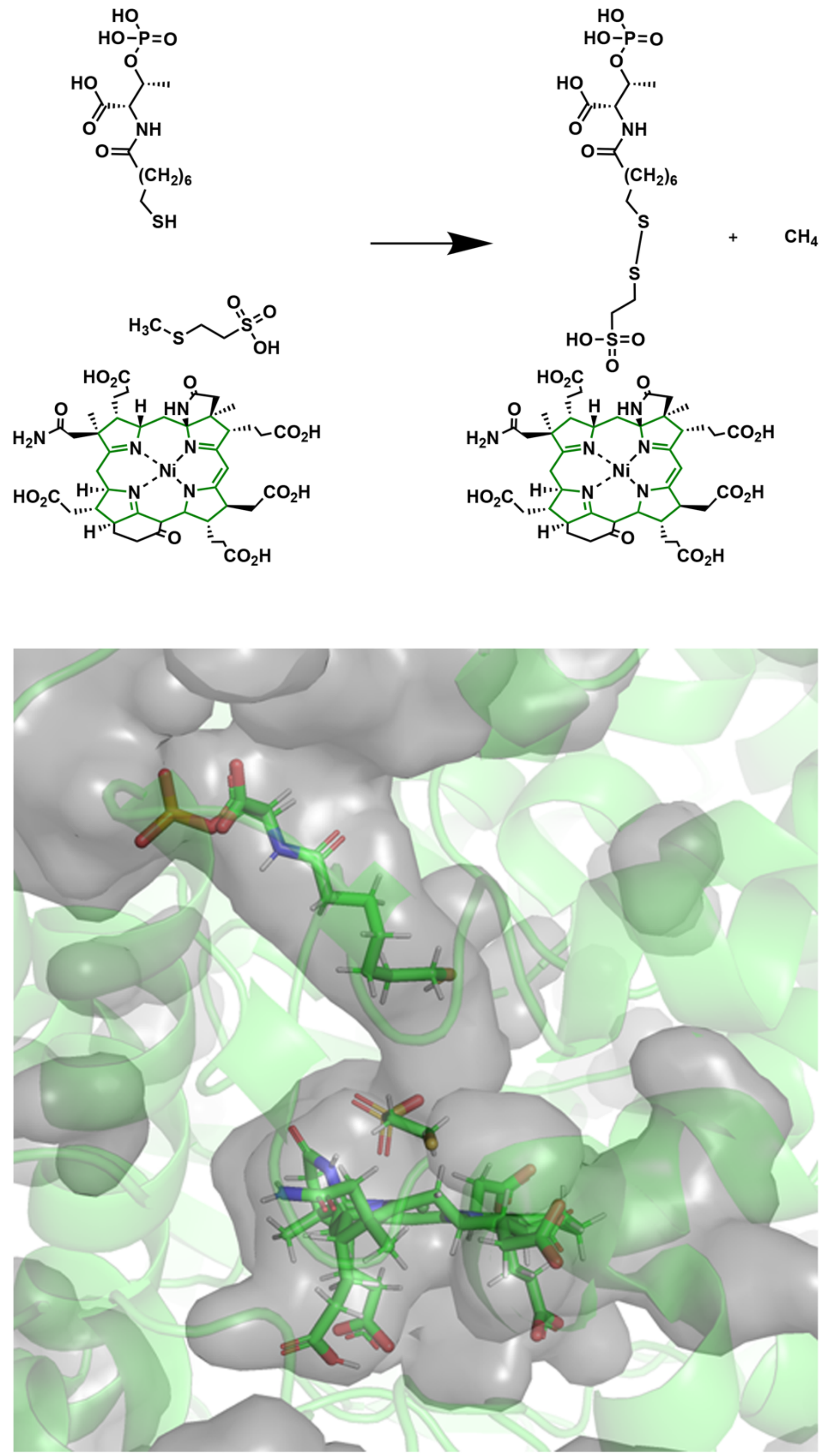

**Extended Data Figure 1|** (Top) Overall Reaction catalyzed by MCR employing the Ni(I)-Cofactor F430. (Bottom) Illustration of the 50Å substrate tunnel of MCR leading to the active site catalytic platform supported by the Ni-Cofactor F430. Note that HSCoM is modelled as only HSCoM has been observed in the active site of MCR in crystal structures.

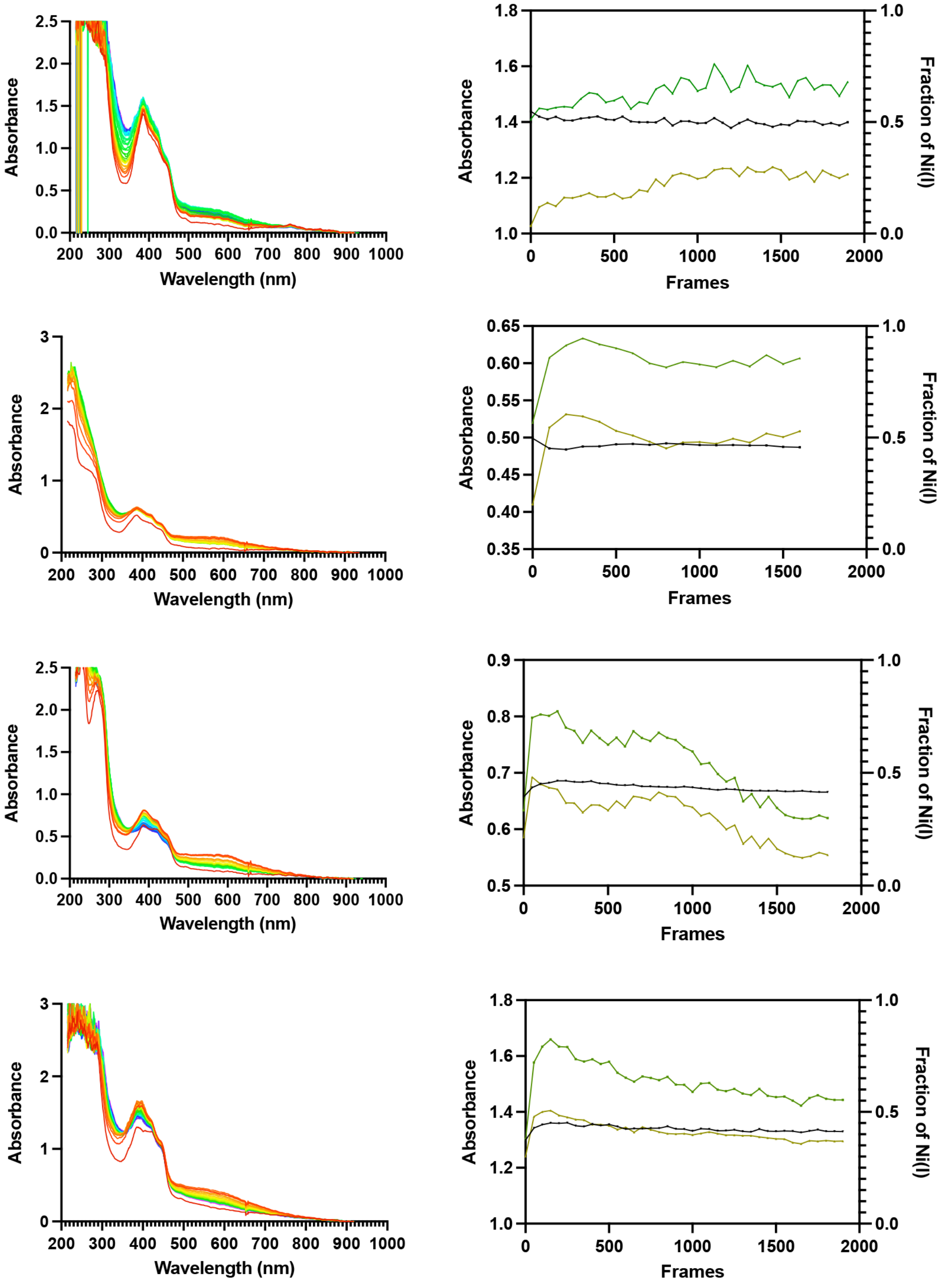


**Extended Data Figure 2| Microspectroscopic analysis of various Ni(I) MCR crystals.** A rainbow color scheme tracks the frames collected during data collection (Red, 0 frames, to Blue, 1800 or 3600 frames) (Inset) Ni(I) percentage (Black line) utilizing a multiwavelength calculation consisting of absorbance values of 385 nm (Green Line) and 420 nm (Yellow Line).

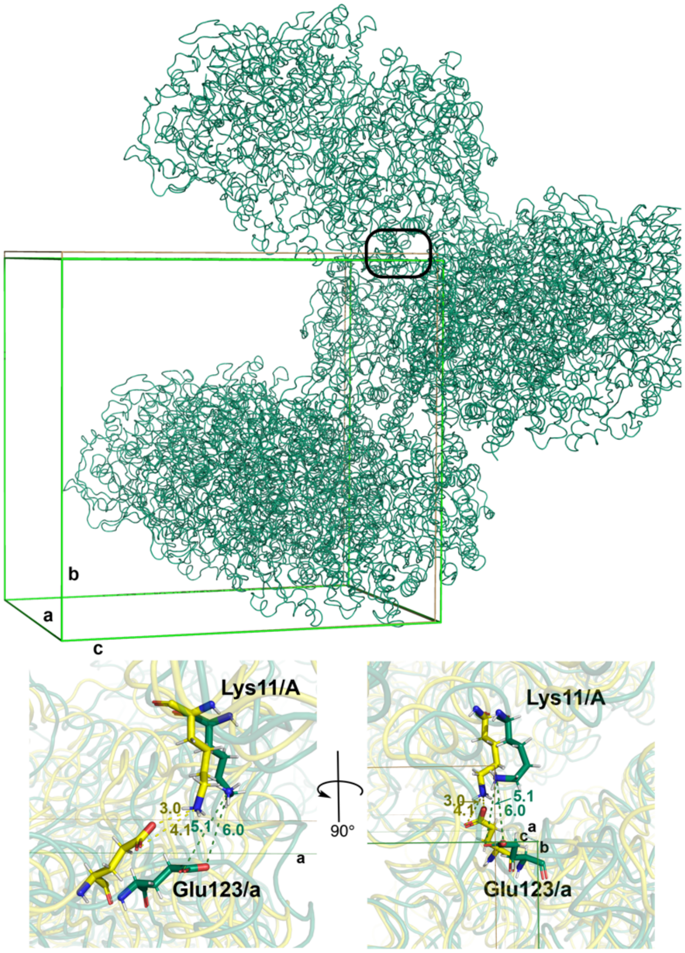


**Extended Data Figure 3|** Unit cell packing analysis reveals critical salt bridge responsible for unit cell differences. **A)** Unit cell packing of MCR where the novel Ni(I) unit cell is illustrated in green and the Ni(II) unit cell is shown in yellow. The Ni(I)/Ni(II) ensemble, shown in green, orients into space group $P12_11$ with the unit cell of a = 83.2 Å, b=118.2 Å, and c=124.5 Å, while the Ni(II)-$MCR_{red1\text{-}silent}$ orients with the same space group, but with unit cell dimensions a = 83.01 Å, b=119.39 Å, and c=123.39 Å. **B)** Ionic interaction leading to the shortening of axis b and lengthening of axis c along the α/α' crystal interface.

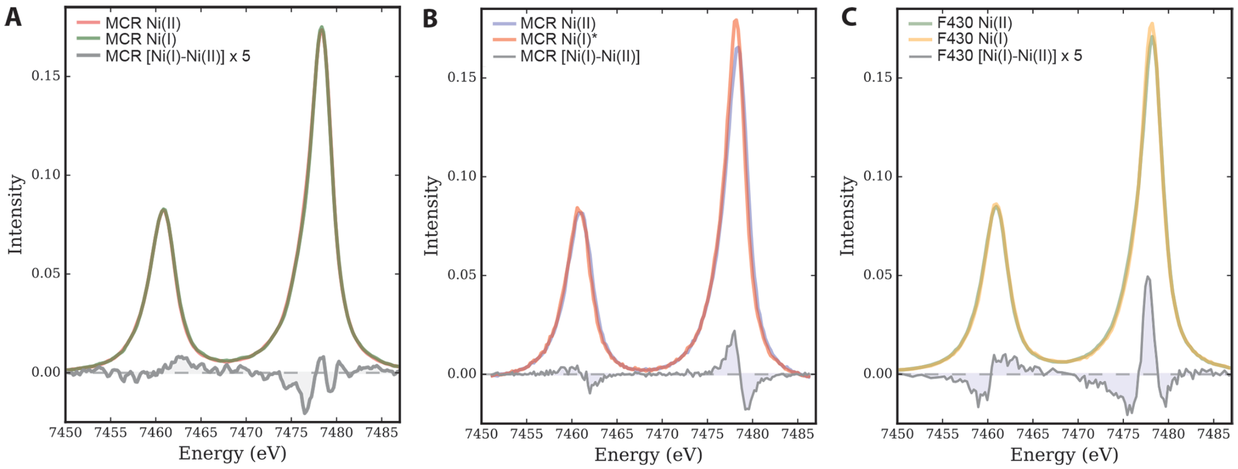


**D**

| Sample | FWHM (eV) | ΔFWHM (eV) |
|---|---|---|
| MCR Ni(I) crystals (XFEL) | 3.20 | |
| MCR Ni(II) crystals (XFEL) | 3.32 | 0.12 |
| MCR Ni(I)* solution (SR) | 2.78 | |
| MCR Ni(II) solution (SR) | 3.07 | 0.29 |
| F430 Ni(I) (SR) | 2.95 | |
| F430 Ni(II) (SR) | 3.16 | 0.21 |

**Extended Data Figure 4**| Ni Kα X-ray emission spectra of MCR samples. **(A)** Ni Kα X-ray emission spectra of crystals collected in tandem with SFX resulting in the Ni(**I**) containing 1.50Å ensemble structure and the 1.45Å Ni(**II**)-inactive structure in green and magenta, respectively. Difference spectrum is shown below in grey. (**B**) Ni Kα X-ray emission spectra of frozen solutions of Ni(**II**) $MCR_{red1\text{-}silent}$ (blue), Ni(**I**) $MCR_{red1}$ (magenta; generated by subtracting 31% of $MCR_{red1\text{-}silent}$), and difference spectra (grey). (**C**) Ni Kα X-ray emission spectra of frozen solutions of the isolated F430 cofactor in the Ni(**II**) and Ni(**I**) form in green and yellow, respectively, and the difference spectrum below in grey (**D**) Tabulated overview of full width half max (FWHM) and difference in FWHM.

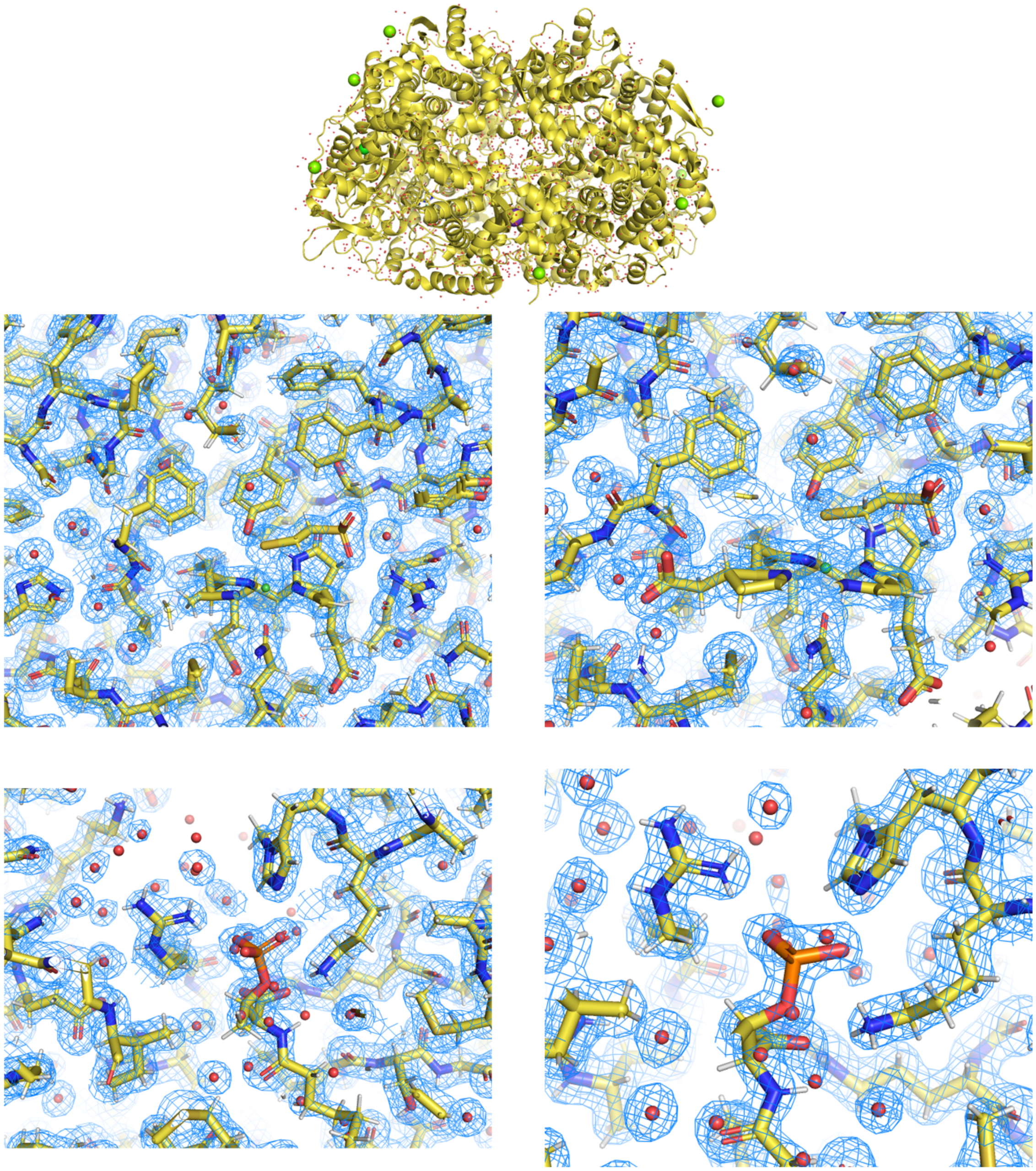

**Extended Data Figure 5| (Top**) Oxidized XFEL structure after *in situ* air oxidation of the Ni(I) MCR sample solved at 1.45Å resolution (pdb:xxxx). **(Middle)** Active site view of both protomers of the assymetric subunit showing the Ni coordinated lower axial Gln147α and CoM thiolate. **(Bottom)** CoB substrate binding tunnel illustrating the "locked-in" nature of the Ni(II) inactive MCR. Mesh is represented as the 2Fo-Fc map at 1.0σ. Refer to Supplemental Table 1 for crystallographic statistics related to the Ni(II) protein.

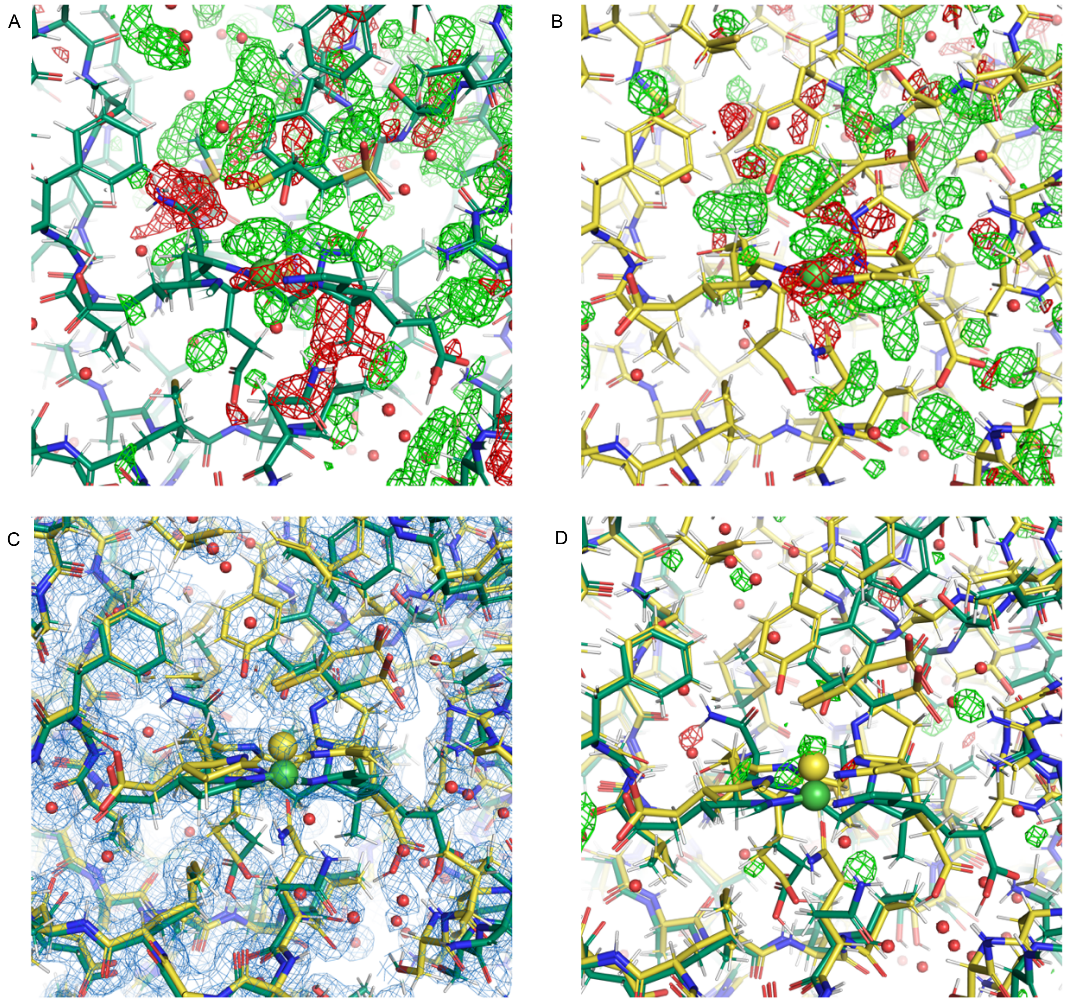


**Extended Data Figure 6|** Difference map illustrating the need for an ensemble model for density fitting. **A)** Fo-Fc map of a refined Ni(I)MCR model with a 1.0 occupancy leading to large areas of positive and negative density at a contour of 3**σ** showing the need for a Ni(**II**) model. **B)** Fo-Fc map at 3**σ** of a refined Ni(**II**)MCR model illustrating the need for a Ni(**I**) model addition. **C)** Active site architecture displaying the models of the Ni(**I**) (green) and Ni(**II**) (yellow) MCR accompanied by the 2Fo-Fc map at a contour of 1σ. **D)** Fo-Fo difference map of both Ni(**I**) and Ni(**II**) models demonstrating convergence of the model with the data collected. Refer to Supplemental Table 1 for crystallographic statistics related to the XFEL ensemble structures of the Ni(**I**) (designated as ALTLOCA) (and Ni(**II**) (ALTLOCW) proteins (pdb:xxxx).

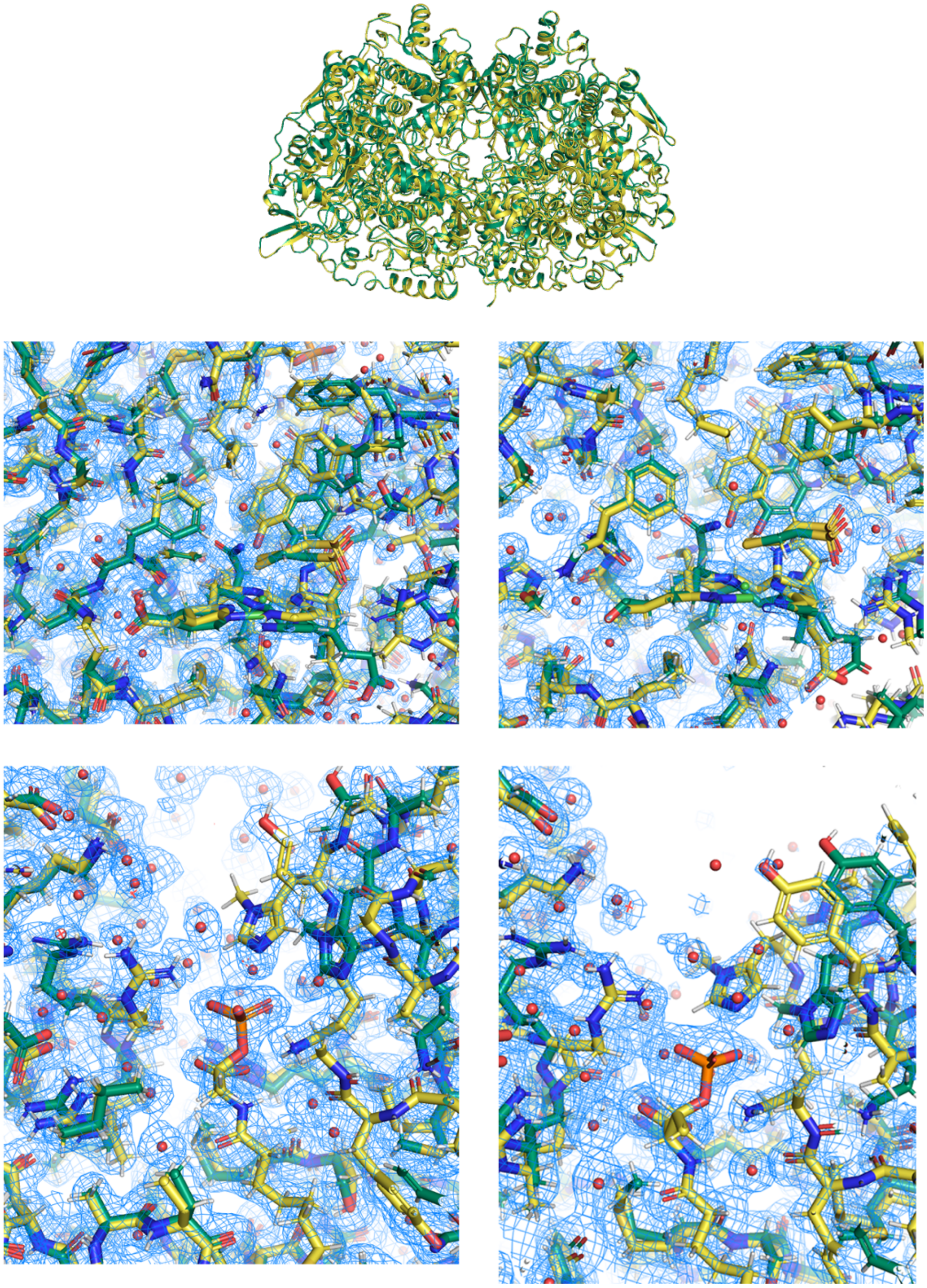

**Extended Figure 7.|** (Top) Cryogenic capillary XRD structure of the Ni(I) MCR sample solved at 1.55Å resolution where the Ni(I) conformation is colored green and the Ni(II) conformation is colored yellow (pdb:xxxx). (Middle) Active site view of both protomers of the assymetric subunit showing a removal of the Ni coordinated lower axial Gln147α and CoM thiolate. (Bottom) CoB substrate binding tunnel illustrating an opening of the substrate tunnel in the Ni(I) conformation. Mesh is represented as the 2Fo-Fc map at 1.0σ. Refer to Supplemental Table 1 for crystallographic statistics related to this capillary XRD structure of the Ni(I) protein.

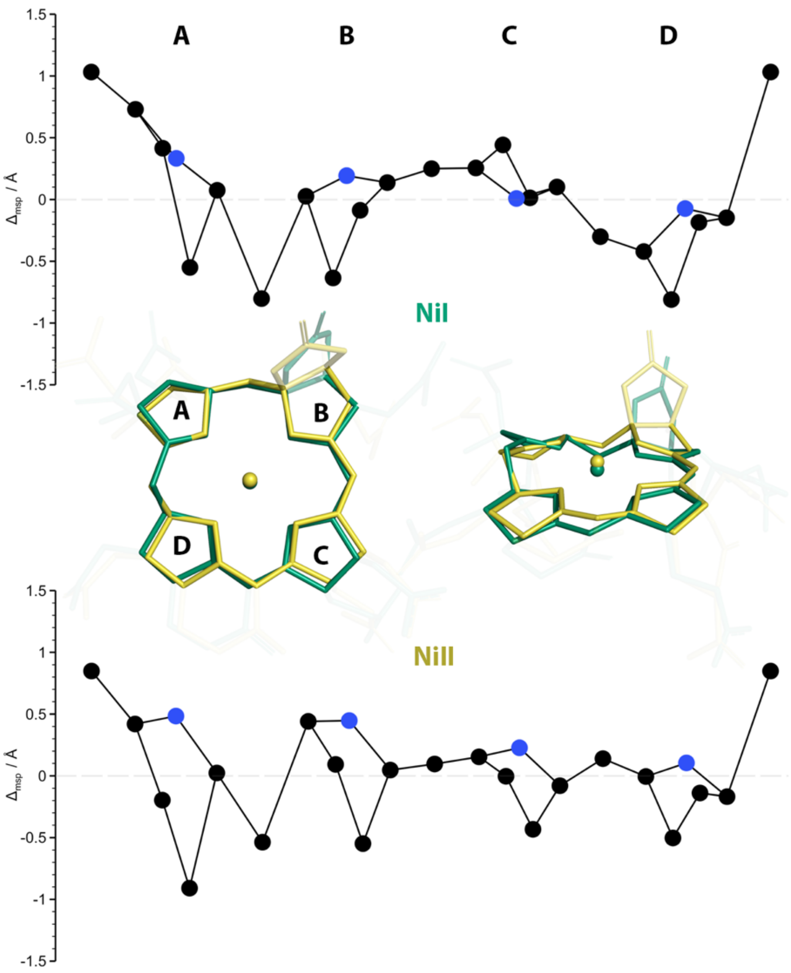


**Extended Data Figure 8|** Porphystruct displacement diagrams illustrate the doming effect of the Ni(II) structure where the dotted line represents the mean plane of the corphin, black data points represent the carbon molecules of the corphin, and blue data points represent the Ni coordinating nitrogens.

A

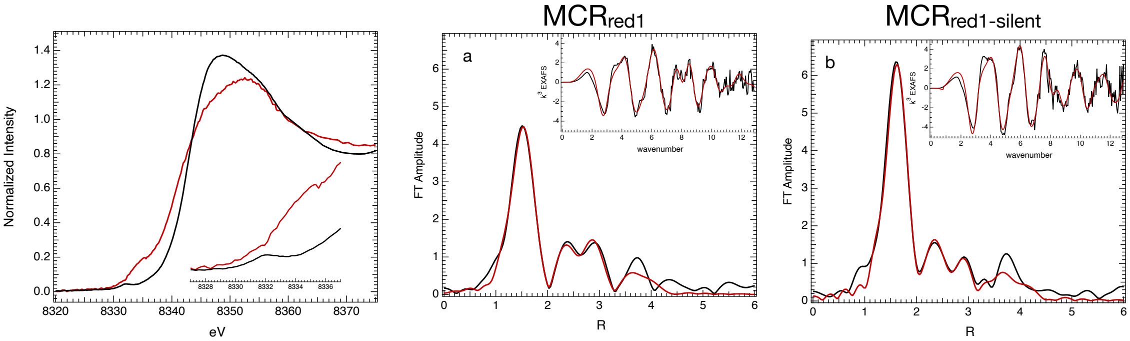


EXAFS least-squares fitting parameters.

| Path | Coordination | R(Å)[a] | $\sigma^2$ (Å$^2$)[b] | ΔE | K range | R-factor | $\chi^2_{red}$ |
|---|---|---|---|---|---|---|---|
| | | | **$MCR_{red1}$** | | | | |
| N | 4 | 2.025 | 476 | -0.04 | 2.5-12.3 | 0.021 | 31.7 |
| O | 1 | 2.157 | 180 | | R range | $N_{var}$ | $N_{idp}$ |
| S | 0.15 | 2.472 | 62 | | 1-4.4 | 17 | 20.9 |
| C | 6 | 3.03 | 606 | | | | |
| N-C | 12 | 3.07 | 312 | | | | |
| C | 3 | 3.36 | 470 | | | | |
| S | 1 | 3.92 | 942 | | | | |
| C-C/N | 12 | 4.38 | 710 | | | | |
| | | | **$MCR_{red1-silent}$** | | | | |
| Path | Coordination | R(Å)[a] | $\sigma^2$ (Å$^2$)[b] | ΔE | K range | R-factor | $\chi^2_{red}$ |
| N | 4 | 2.084 | 347 | 2.5 | 2.5-12.5 | 0.021 | 25.5 |
| O | 1 | 2.185 | 347/ | | R range | $N_{var}$ | $N_{idp}$ |
| S | 1 | 2.434 | 816 | | 1-4.5 | 14 | 22 |
| C | 5 | 3.04 | 755 | | | | |
| N-C | 10 | 3.06 | 1613 | | | | |
| C | 4 | 3.37 | 1160 | | | | |
| N-C | 12 | 4.43 | 942 | | | | |

[a]The estimated standard deviations in interatomic distance is ± 0.02 Å. [b]The $\sigma^2$ values have been multiplied by $10^5$. A "/" denotes that the value was shared with that of the prior path.

B

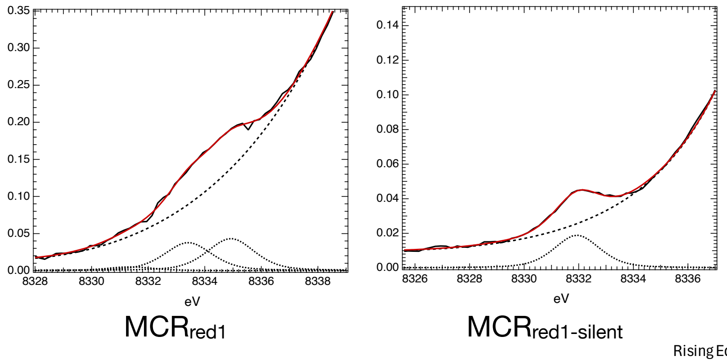


| Sample | Edge Inflection | μE = 0.5 | Pre-edge Position | Pre-edge Intensity | Rising Edge Feature Peak 1 | Rising Edge Feature Peak 2 |
|---|---|---|---|---|---|---|
| **MCR_red1_silent** | 8342.5 | 8341.5 | 8331.9 | 0.047 | n/a | n/a |
| **MCR_red1** | 8340.9 | 8340.0 | 8331.4 | 0.013 | 8333.4 | 8334.9 |

*calibrated to 8331.6 eV, FWHM = 1.97 eV

**Extended Data Figure 9| Ni K-edge XAS of MCR crystal slurries**. **(A. Top)** *Inset*: expanded pre-edge region. $MCR_{red1}$ (—), $MCR_{red1-silient}$(—). (**A. Bottom**) Ni K-edge EXAFS of $MCR_{red1}$ (a) and $MCR_{red1-silent}$ (b) crystals slurries. The edge positions measured both as the maximum of the first derivative (edge inflection) and at μE = 0.5 are presented. **(B)** The pre-edge position, intensity, and positions of the peak in the rising edge of the Ni(**I**) species were determined from pseudo-Voigt Peak deconvolution. Fit values are presented in table below.

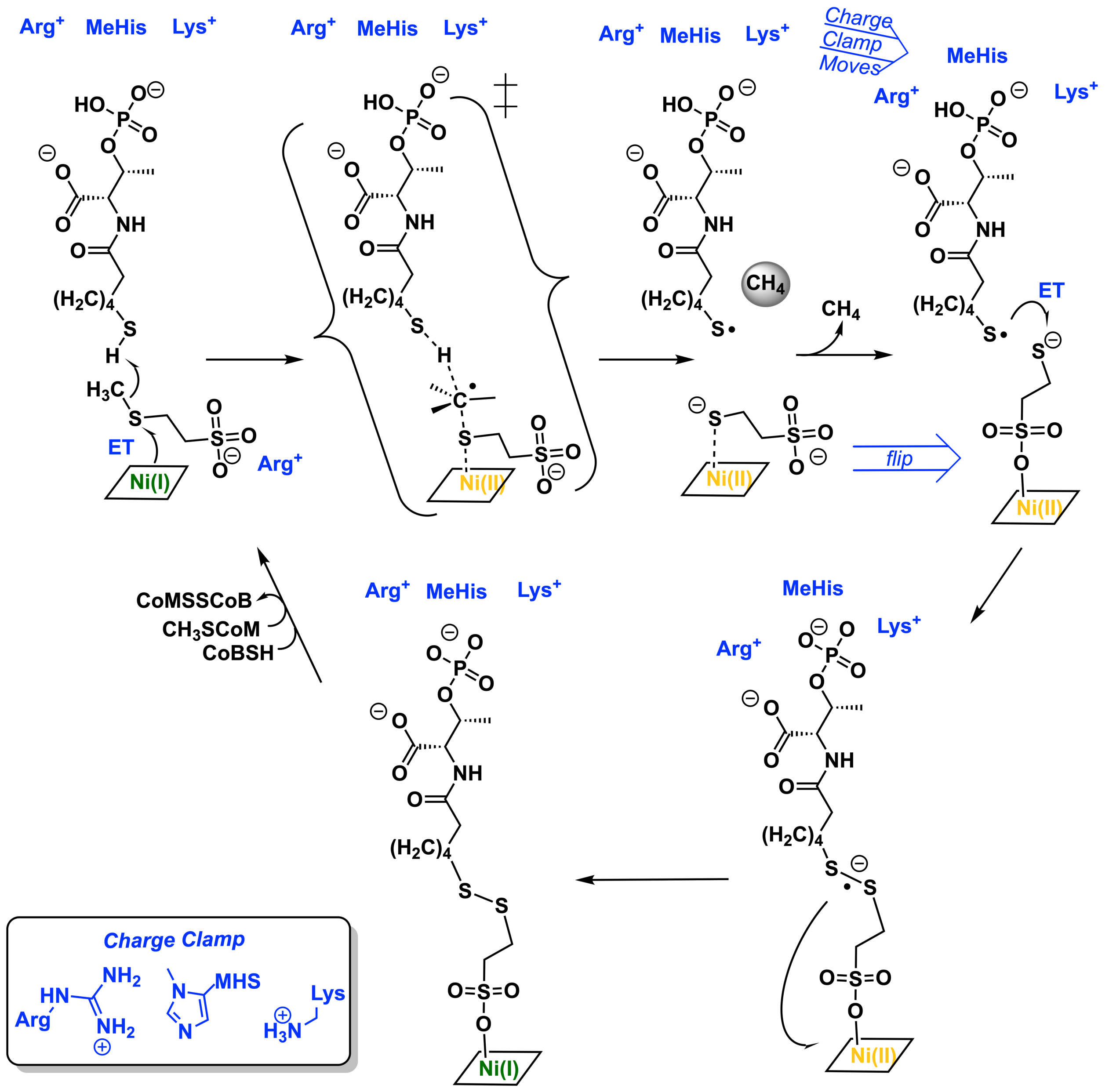


**Extended Data Figure 10|** Proposed mechanism of MCR based on the Ni(I) MCR structure.
